# Capturing α-synuclein polymorphism with AlphaFold 3 and rivals

**DOI:** 10.64898/2025.12.05.692529

**Authors:** Rebecca M. Price, Ben LaCourse, Aderik Voorspoels, Luc G. Elliott, Alana Maerivoet, Adam J. Simpkin, Shahram Mesdaghi, Alexandra Stafford-Dodsworth, Jillian Madine, Daniel J. Rigden

## Abstract

Amyloid fibrils formed by a single protein can adopt many distinct folds, or polymorphs, which differ between diseases and shape the surfaces available to therapeutics and diagnostics. Whether deep-learning structure prediction can capture this diversity is unclear, and the field lacks an objective definition of a polymorph from which to judge this. Here we cluster the α-synuclein structures in the Protein Data Bank by TM-score, giving an objective reference set of known polymorphs. Against this reference we assess AlphaFold 3, Boltz-2 and OpenFold 3 run through ABCFold. All three produced fibril-like assemblies, though they differed in which polymorphs they reached. With templates included as default the methods over-represented the dominant fold. When untemplated the methods produced a substantial number of models matching no known structure, a subset of which have the stacking geometry of genuine fibrils. Supplying a polymorph as a template recovered 13 of 16 target folds, but only when alignment depth and the number of modelled chains were varied together. These results establish that current methods sample real α-synuclein structural space and can be steered through it, and provide a framework for assessing prediction methods on amyloid targets.

## Introduction

Amyloid fibrils are highly ordered protein assemblies formed by the aggregation of amyloidogenic proteins into highly stable structures that normally possess a cross-β structure. The cross-β structure consists of β-strands running perpendicular to the fibril axis, stacked into β-sheets held together by inter-strand hydrogen bonds that run parallel to the axis (Nelson *et al*., 2005; Sawaya *et al*., 2007a). Their formation is most commonly associated with protein misfolding (Otzen and Riek, 2019), whereby the monomeric soluble protein must first transition into a structure that allows for aggregation. Subsequent steps involve the formation of higher order oligomers, protofibrils and finally mature fibrils (Verma, Vats and Taneja, 2015). Accumulation of amyloid fibrils, both intracellularly and extracellularly, is linked to neurodegenerative disease (LaFerla, Green and Oddo, 2007; Rambaran and Serpell, 2008). In Parkinson’s disease (PD), fibrils are formed from misfolded α-synuclein and are thought to contribute to the disease pathogenesis (Chiti and Dobson, 2006).

High-resolution structures of amyloid fibrils emerged decades after the first globular protein structures were solved in the 1950s and 60s (Kendrew *et al*., 1958; Perutz *et al*., 1960), reflecting both the complex intrinsic properties of the fibrils and limitations of earlier structural methods. Amyloid fibrils are insoluble, non-crystalline, and heterogeneous, making them incompatible with traditional X-ray crystallography and solution NMR. Early low-resolution models appeared in the 2000s (Petkova *et al*., 2002; Sawaya *et al*., 2007b), but a breakthrough came with advances in cryo-electron microscopy (cryo-EM), including direct electron detectors (Li *et al*., 2013) and improved helical reconstruction algorithms (He and Scheres, 2017a), producing high-resolution structures of amyloid-β, tau, α-synuclein, and immunoglobulin light chain amyloid fibrils (Fitzpatrick *et al*., 2017; Gremer *et al*., 2017; Li *et al*., 2018a; Radamaker *et al*., 2019). Cryo-EM’s compatibility with the elongated, helical morphology of fibrils has led to a substantial increase in the number of characterised amyloid structures.

Despite such advances, amyloid fibrils remain exceptionally difficult to model computationally. As fibril formation arises from misfolding, the architecture does not reflect an evolutionarily optimised fold, meaning sequence-derived co-variation and evolutionary couplings, key signals leveraged by modern deep-learning predictors such as AlphaFold 2 (AF2) are weak or absent (Jumper *et al*., 2021). In contrast, functional amyloids for which amyloid is their native conformation, have been plausibly modelled using covariance based structure prediction methods (Tian *et al*., 2015; Sønderby *et al*., 2022; Mesdaghi *et al*., 2023). Adding to the difficulties in pathological amyloid structure prediction, a single polypeptide can form multiple, structurally distinct polymorphs with differing protofilament and β-sheet arrangements (Paravastu *et al*., 2008; Tycko, 2015). The lack of standardised nomenclature for these polymorphs further complicates cross-study comparison (Eisenberg and Sawaya, 2017).

Recent advances in deep-learning-based structure prediction offer new opportunities for studying amyloids. While AF2 excels at predicting monomeric, globular protein structures, it struggled with aggregation-prone or misfolded states (Pinheiro, Santos and Ventura, 2021) and initially required concatenated sequences with artificial chain breaks for multimer prediction (Jumper *et al*., 2021; Bæk and Kepp, 2022). In contrast, AlphaFold 3 (AF3) and similar methods improve multimer predictions and inter-chain contacts (Abramson et al., 2024). Their diffusion-based structure modules also generate multiple sampled structures rather than a single refined prediction, although recovery of genuinely alternative conformations remains limited without manipulation of the input multiple sequence alignment (MSA) (Chakravarty *et al*., 2024)

Here, we assess how well AF3 and two other deep-learning methods, Boltz-2 and OpenFold 3 (OF3), predict α-synuclein fibril structures. We first cluster the experimentally determined α-synuclein structures in the protein data bank (PDB) by TM-score, giving an objective definition of the known polymorphs. Using this reference, we ask which polymorphs each method reproduces by default, whether models resemble structures released after each method’s training cut-off or represent plausible novel folds, and how far each method can be steered towards a specific polymorph using templates.

## Methods

### Structure prediction

Initial AF2 models were generated through ColabFold (Mirdita *et al*., 2022) with default parameters unless otherwise stated, using the α-synuclein sequence from UniProt entry P37840 (UniProt Consortium, 2023). For custom templating, the MSA was reduced to a single sequence and PDB structure 6CU7 was supplied as a custom template. Initial AF3 models were generated through the AlphaFold Server (https://alphafoldserver.com/) with default parameters (Abramson *et al*., 2024).

All further models were generated with ABCFold (v1.1.6; (Elliott, Simpkin and Rigden, 2025)) using AF3 (v3.0.1), Boltz-2 (v2.2.1; (Passaro *et al*., 2025)) and OpenFold 3 (v0.4.3; (The OpenFold3 Team, 2026)). The fibril-forming core of α-synuclein (residues 37-97) was modelled as 1-15 chains, with five models per method for each oligomer size. Full-depth MSAs were generated with MMseqs2 (Steinegger and Söding, 2017), and each oligomer size was run with and without ABCFold’s template search of the PDB.

For template-guided modelling, custom templates were used for each of the 16 reference polymorphs. Residues were mapped using the author numbering of each template, as the resolved range differs between structures. Sequence identity at each mapped position was checked against the canonical α-synuclein sequence; the only mismatches were known point mutations (e.g. H50Q, G51D, A53T). Each template was supplied to AF3, Boltz-2 and OF3, with residues 37-97 modelled as 1-8 chains. MSA depth was varied at 1, 2, 4, 8, 16, 32 and the full MSA (71 sequences), with shallower MSAs taken as the first n sequences of the full alignment. MSAs were supplied directly as input. Five models were generated per condition.

### α-Synuclein reference set and clustering

All structures in the PDB (Berman, Henrick and Nakamura, 2003) associated with UniProt P37840 were downloaded (accessed 08/2026). Chains were kept if they shared ≥85% identity with the α-synuclein sequence, allowing for point mutations, and had at least 15 resolved residues. Structures with three or more such chains were classed as fibrils, and the 18 structures with only one or two chains were inspected individually. Within each structure, chains were grouped by complete-linkage hierarchical clustering of pairwise Cα RMSD (Biopython; (Cock *et al*., 2009) 1.0 Å cutoff, ≥10 shared residues), and one chain was kept per group, giving 224 chains from 196 structures.

Pairwise TM-scores between the 224 chains were calculated with Foldseek (van Kempen et al., 2024), converted to distances (1 − TM-score), and clustered by average linkage using a custom Python script, with clusters defined at a distance cutoff of 0.5. The medoid of each cluster was taken as the chain with the highest mean TM-score to all other members. Chains without a cluster partner at this cutoff were classed as singletons and numbered 9-16. Principal component analysis (10 components) was performed on the symmetrised TM-score matrix using scikit-learn (Pedregosa *et al*., 2011). To define conserved cores, chains were superposed on their cluster medoid by iterative fitting on Cα atoms within 2 Å, repeated until the fitted set converged.

### Model analysis

Initial AF3 models were compared with the PDB using Foldseek (van Kempen *et al*., 2024). For ABCFold models, backbone clashes were identified with SciPy cKDTree (Virtanen *et al*., 2020) as any pair of backbone atoms (N, Cα, C, O) closer than 2.0 Å, within or between chains, excluding adjacent residues in the same chain. Models containing a clash were removed. Chains within each remaining model were grouped by Cα RMSD (1.0 Å, complete linkage) as for the PDB structures, and one chain per group was kept for further analysis.

Models were classified as fibril-like if they contained at least two chains, at least 80% stacked residues and at least 25% β-strand residues. Stacked residues were defined as those whose Cα lay within 4.4-5.5 Å of the Cα of the equivalent residue in another chain, a range around the 4.8 Å inter-strand spacing of a cross-β fibril. β-strand content (DSSP codes E and B) was assigned with DSSP (Kabsch and Sander, 1983) on the full assembly, since cross-β structure depends on inter-chain hydrogen bonds. Thresholds were set by applying the same measures to α-synuclein fibril structures in the PDB with three or more chains. Helical rise and twist were calculated for fibril-like models and PDB structures with at least four stacked chains, from a Kabsch superposition (Kabsch, 1976) of the first and last chain, ordered along the fibril axis by PCA, divided by the number of chain steps. Twist is signed, with negative values left-handed, and structures with a twist magnitude below 0.5° were classed as near-planar.

Each chain was compared with the 224 PDB chains using Foldseek in TM-align mode and assigned to the cluster of its closest match if the TM-score was 0.5 or higher. Fibril-like chains without a match were classed as novel, and the remainder as non-fibril. Novel chains were then compared all-against-all in the same way and clustered as for the PDB structures. To compare with training data, PDB chains were labelled by release date relative to each method’s training cut-off (AF3 and OF3, 30 September 2021; Boltz-2, 1 June 2023). For models matching a structure released after the cut-off of the method that produced them, the highest TM-score to any pre-cut-off structure was also recorded.

For template-guided models, each chain was scored by TM-score to its template, and the highest value across the five models of each condition was taken. A fold was counted as recovered at a TM-score of 0.5 or higher.

Data were visualised using matplotlib (https://matplotlib.org/) and seaborn (https://seaborn.pydata.org/), and structures were visualised and figures generated in PyMOL (https://www.pymol.org/).

## Results

### AlphaFold 3, but not AlphaFold 2, produces fibril-like α-synuclein assemblies

To investigate the ability of AF2 and AF3 to predict fibril assemblies, trimeric models were generated for both full-length and truncated α-synuclein (37-97). Initial AF2 predictions failed to produce fibril assemblies, often resulting in implausible disordered or α-helical structures exhibiting multiple chain clashes and low predicted local distance difference test (pLDDT) scores. In the case of modelling full length α-synuclein, a direct assembly of the monomeric structure is produced (Figure 1A), and a similar α-helix rich structure is seen for truncated α-synuclein (Figure 1B). Although pathological amyloid arises as a result of a misfolding, the inability to produce a fibril model is perhaps still surprising when considering the number of fibril structures of α-synuclein in the PDB compared to only a handful of monomeric structures.

**Figure 1.**
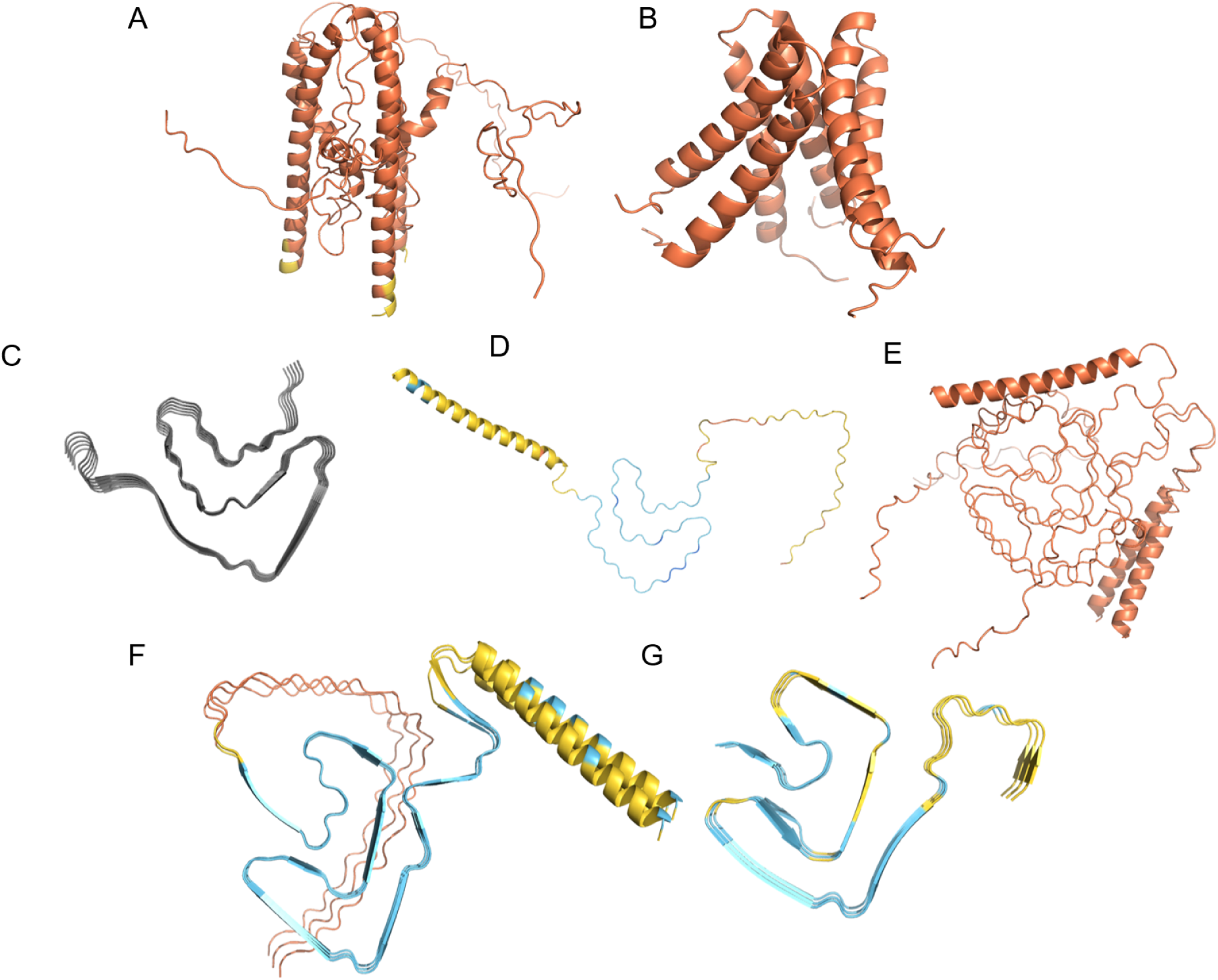
α-synuclein trimers predicted by AF2 and AF3. A) Full-length and B) truncated (37-97) α-synuclein trimers predicted by AF2 with default parameters. C) PDB structure 6CU7. D) Full-length α-synuclein monomer and E) trimer predicted by AF2 using 6CU7 as a template and a single sequence input. F) Full-length and G) truncated α-synuclein trimers predicted by AF3. Predicted structures are coloured by pLDDT (dark blue > 90, light blue 70-90, yellow 50-70, orange < 50)

AF2 can be guided to produce a fibril-like fold for α-synuclein, but this requires changes from default parameters. Reducing the MSA depth to a single sequence input, in addition to custom templating with a PDB fibril structure (6CU7; Figure 1C), results in a fibril-like fold for monomeric α-synuclein corresponding to the given template (Figure 1D). Templating without the removal of the MSA is not sufficient to yield a fibril structure. However, when the assembly is expanded to a trimer, inter-chain clashes again dominate, preventing the formation of a plausible fibrillar structure (Figure 1E). Due to these issues, subsequent modelling focused on AF3.

In contrast, when a trimeric model of full length α-synuclein is modelled by AF3 with default parameters, including templates and full depth MSA, a confident fibril structure is produced (Figure 1F). All five models produced for α-synuclein, both full length and truncated, resemble fibril structures, with the truncated models showing slight differences in morphology whilst still resembling the same broad fold (Figure 1G).

To evaluate the similarity of the AF3-predicted assemblies to experimental structures, the analysis focused on truncated α-synuclein models spanning residues 37-97, since the terminal regions are often unresolved in experiments. Screening against the PDB using Foldseek (van Kempen et al., 2024) returned several hits for each model, matching PDB entries 7NCK (Lövestam *et al*., 2021) and 8BQV (Yang *et al*., 2023), which have the same general fold but do possess some differences (Table 1).

**Table 1.** Top Foldseek hit in the PDB for each AF3 model of truncated α-synuclein. Trimers of residues 37-97 were predicted with the AlphaFold Server using default parameters, including templates and full-depth MSA. For each of the five models, the top-ranked PDB hit, its Foldseek probability and E-value, and the mean pLDDT of the model are given.

| Model | PDB hit | Probability | E-value | Mean pLDDT |
| --- | --- | --- | --- | --- |
| asyn_37_97_model_0 | 7nck | 1 | 2.28E-05 | 67.9 |
| asyn_37_97_model_1 | 7nck | 1 | 4.78E-05 | 68.15 |
| asyn_37_97_model_2 | 8bqv | 1 | 9.50E-05 | 65.98 |
| asyn_37_97_model_3 | 8bqv | 1 | 4.98E-04 | 66.63 |
| asyn_37_97_model_4 | 8bqv | 1 | 7.14E-05 | 65.54 |

Model_0 has the lowest e-value and therefore produced the most significant hit, 7NCK, an α-synuclein filament seeded *in vitro* by filaments purified from a multiple system atrophy patient. Visual inspection revealed a clear resemblance between the two structures, supported by a TM-score of 0.51 (Figure 2) (Zhang and Skolnick, 2005), suggesting the same overall fold but with local differences.

**Figure 2.**
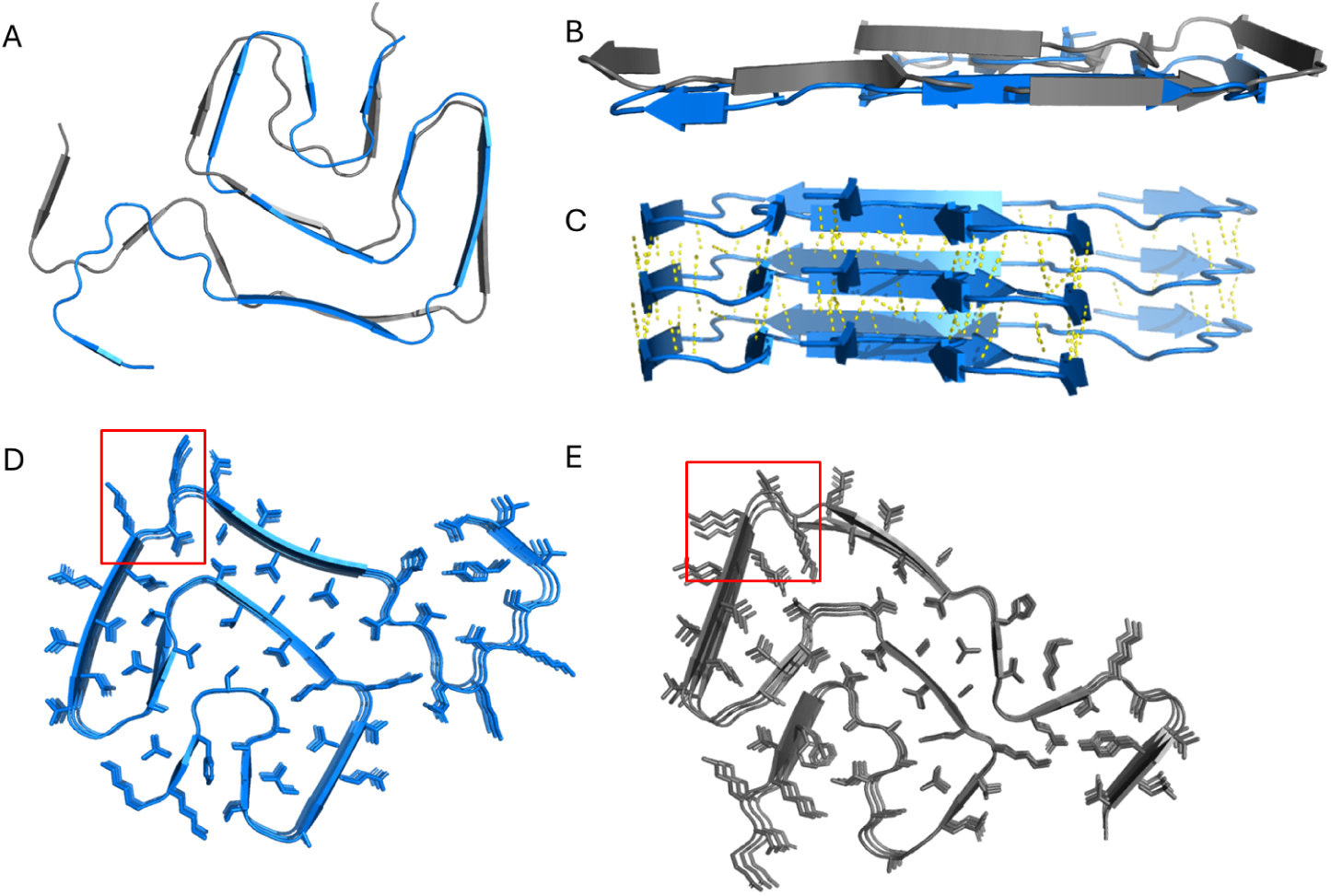
Comparison of the AF3-predicted model_0 of α-synuclein with the experimental fibril structure 7NCK. A) and B) Top-down and side views of a single layer of the AF3 model (blue) aligned to a single layer of 7NCK (grey). The absence of helical pitch in the prediction can be observed in B). C) Hydrogen-bonding network between chains in the AF3 model. Top-down views of D) the AF3 model and E) 7NCK highlighting side-chain orientations. The red box illustrates residues 58-60 where side chain orientation differs between the PDB structure and the AF3 model.

The AF3-predicted model contains eight β-strands per layer, one fewer than the experimental structure, largely driven by differences in both the N- and C-termini, with the core fold largely similar (Figure 2A). Another difference is the lack of helical pitch within the model (Figure 2B). Amyloid fibrils are characterised by three helical parameters. The rise is the translation between successive subunits along the fibril axis, and is around 4.8 Å as it is set by cross-β hydrogen bonding. The twist is the rotation between successive subunits about that axis, and the pitch is the axial distance over a full 360° rotation (Greenwald and Riek, 2010; Taylor and Staniforth, 2022). Experimental α-synuclein fibrils twist by about 1° per subunit, giving pitches of 110 to 280 nm. Here the predicted subunits stack almost without rotation, so the twist is near zero and the pitch correspondingly long. This lowers the alignment scores, because residues that would normally be separated by helical rise remain in the plane, resulting in imperfect superposition despite local structural similarity. Hydrogen bonds between adjacent chains are present in the AF3 model (Figure 2C), in agreement with a cross-β structure, and the bond distances are generally comparable to those observed in 7NCK.

Finally, although the overall backbone architecture of the AF3-predicted model resembles the experimental fibril, there are substantial differences in side-chain orientations. In some regions, side-chain packing agrees well with the experimental model, whereas in others, residues that are buried in the experimental structure appear solvent-exposed in the AF3 model, or vice versa (Figure 2D-E). For example, both lysine 58 and 60 are solvent exposed in the AF3 model, causing threonine 59 to be buried, whereas in 7NCK lysine 58 is buried and stabilised by a nearby glutamic acid (Figure 2D-E, red box). These differences likely arise from subtle shifts in backbone and the absence of the correct helical pitch, which alter the relative positioning of β-strands between layers.

Overall, AF3 was able to capture several key features of amyloid fibril structure whereas AF2 failed to produce comparable structures. Although TM-score is typically benchmarked on globular proteins rather than fibrillar assemblies, the correspondence between the AF3 model and 7NCK suggests that the metric can also capture meaningful similarity between fibril folds. We used α-synuclein as a representative system to further explore the extent to which AF3 and similar prediction methods can both reproduce known fibril forms and predict new, plausible structural variants.

### Exploring deep learning based structure prediction for α-synuclein fibril polymorphs

#### Clustering the PDB

Comparing predicted and experimental fibrils first required a quantitative definition of the experimental polymorphs solved to date. We obtained all PDB entries for α-synuclein (UniProt P37840; accessed 08/2026), removed non-fibrillar structures and short fragments, and extracted one representative chain for each distinct protofilament fold within each structure (see methods). This gave 224 representative chains from 196 structures. Pairwise similarity was scored by TM-score rather than RMSD for two reasons. TM-score down-weights large local deviations, so chains sharing a fibril core are not penalised for differences in flexible or variably resolved termini. It also carries an established, fold-equivalence threshold of 0.5 (Zhang and Skolnick, 2005; Xu and Zhang, 2010). Average-linkage hierarchical clustering at a TM-derived distance of 0.5 separates the 224 chains into eight multi-member clusters and eight singletons (Figure 3, Table 2). As this cutoff applies to the mean linkage between clusters, individual pairs within a cluster can fall below 0.5 while the cluster as a whole remains well supported.

**Figure 3.**
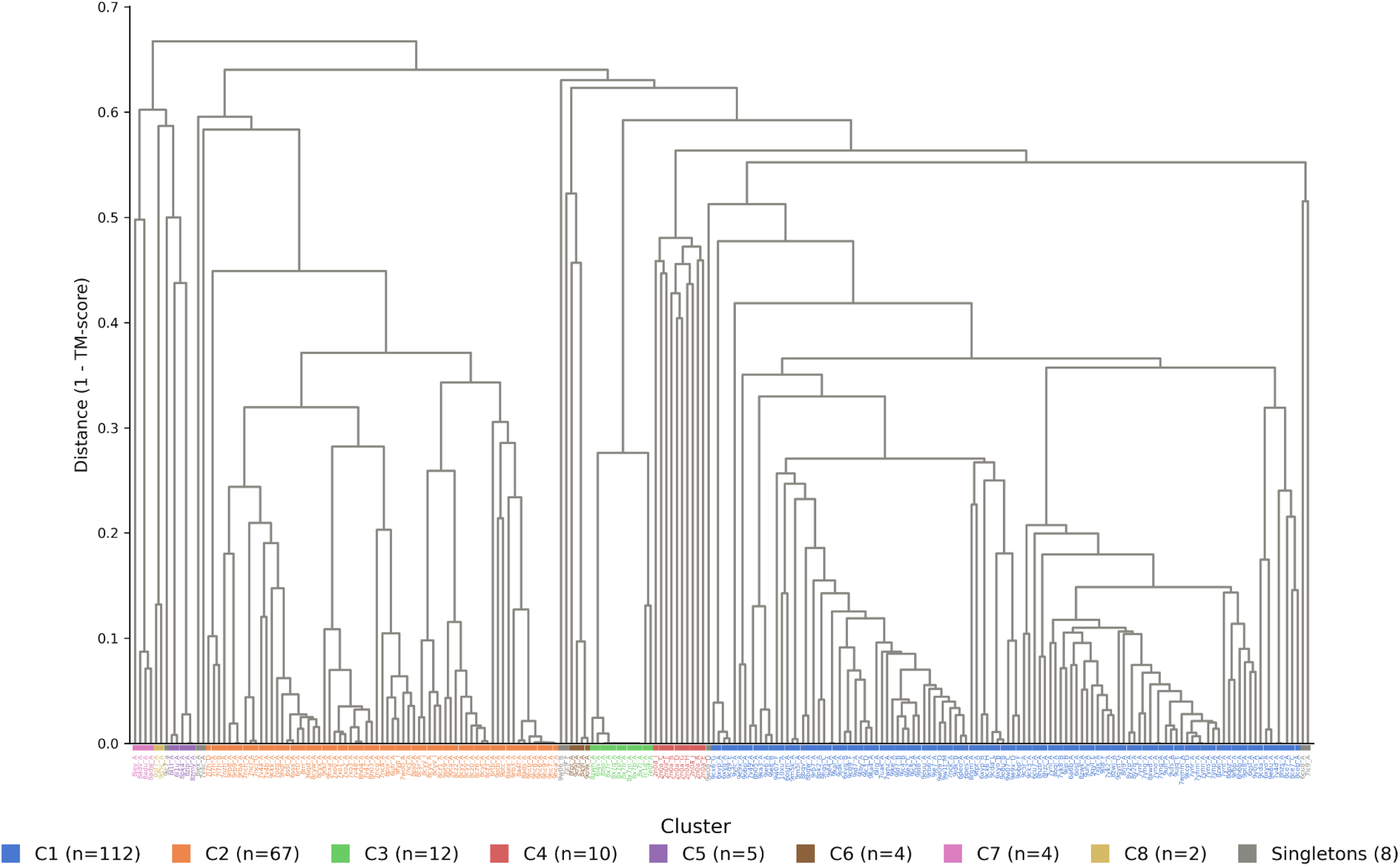
Hierarchical clustering of α-synuclein fibril structures in the PDB. Dendrogram of the 224 representative chains from 196 PDB entries, built by average linkage from all-against-all Foldseek TM-scores converted to distances (1 - TM-score). The tree was cut at a distance of 0.5, giving eight multi-member clusters and eight singletons. Leaf labels give the PDB ID and chain. Labels are coloured by cluster (1-8, as in Table 2), with n in the key giving the number of chains per cluster. Singletons (9-16) are shown in grey.

**Table 2.**
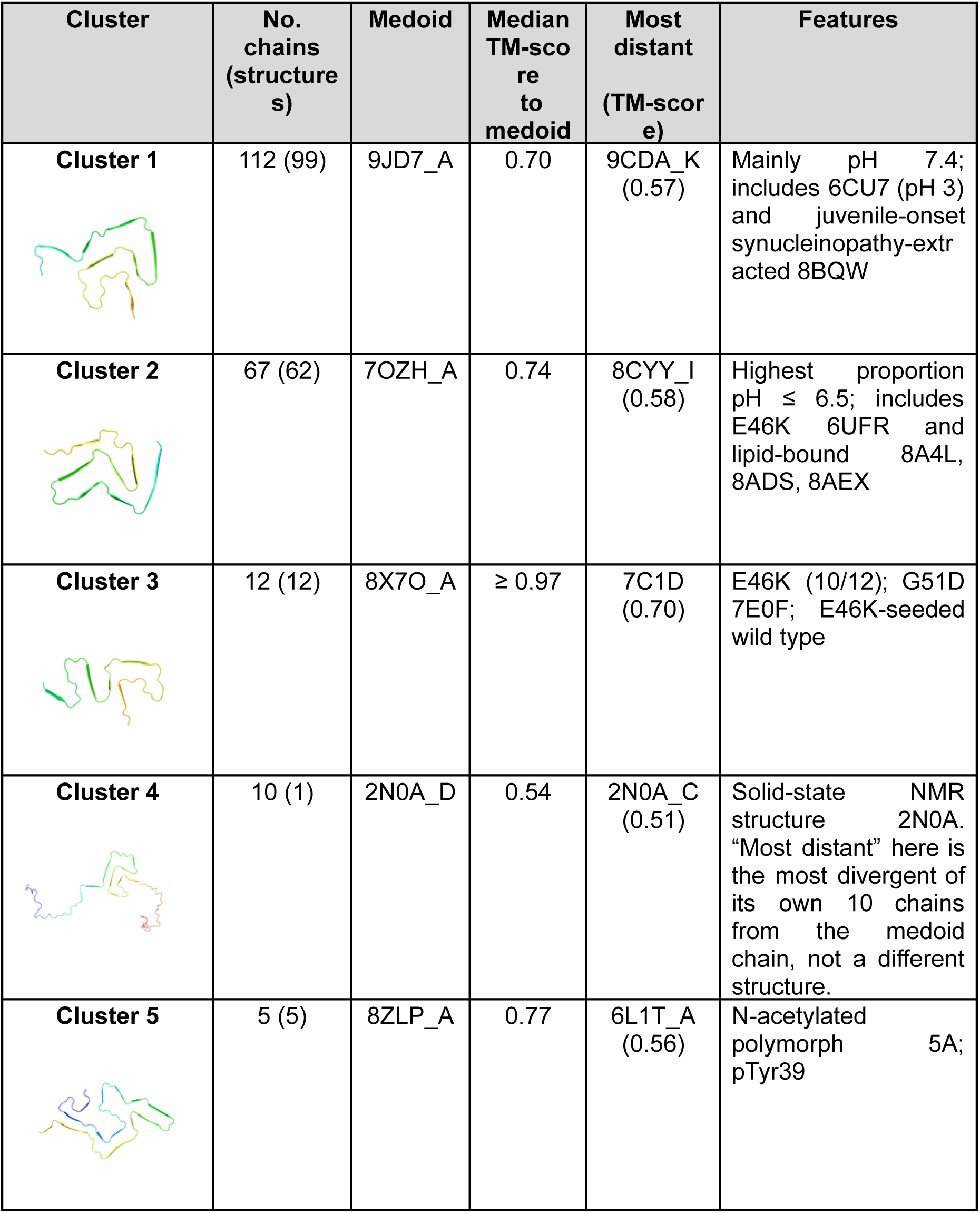
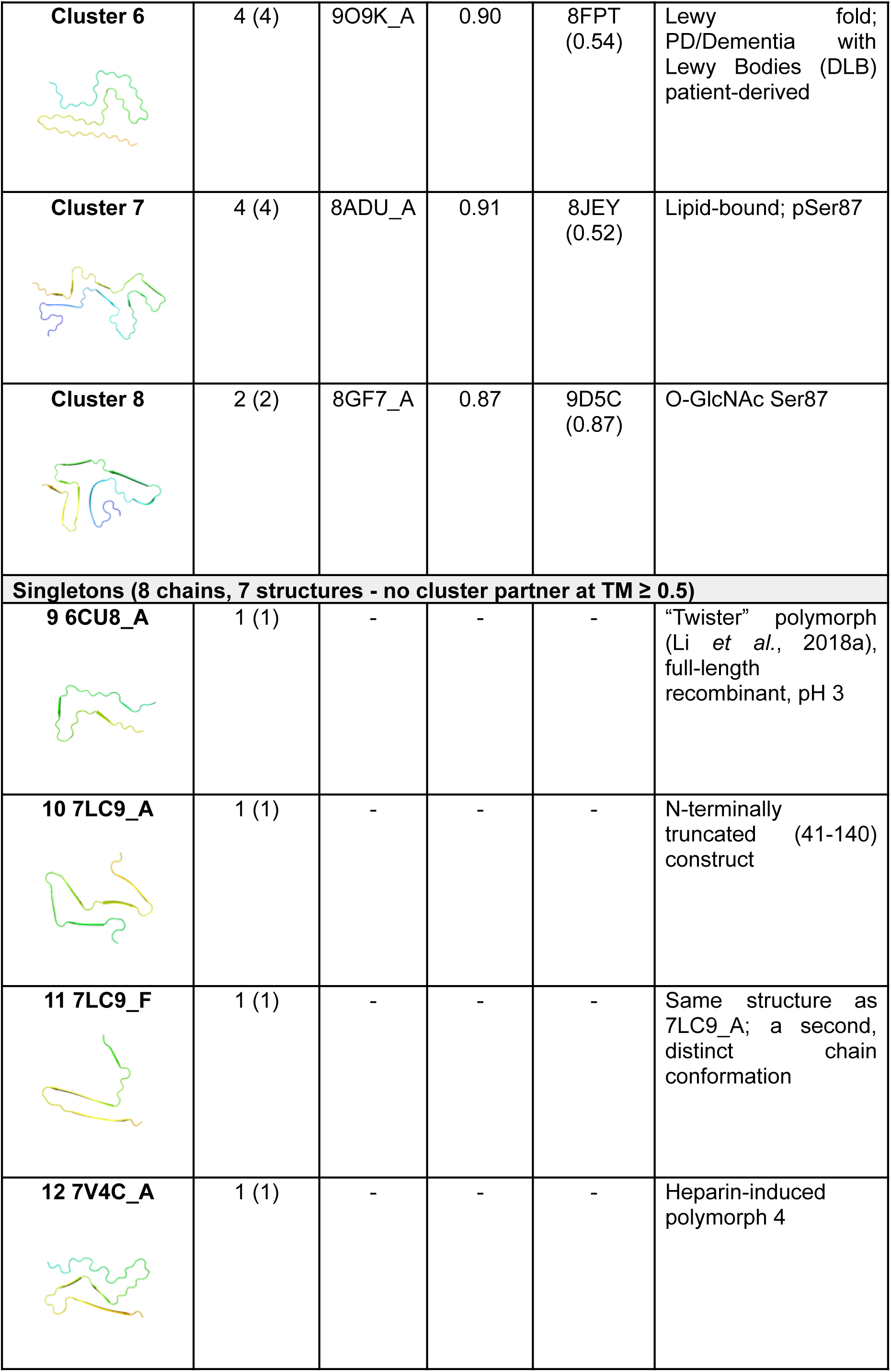
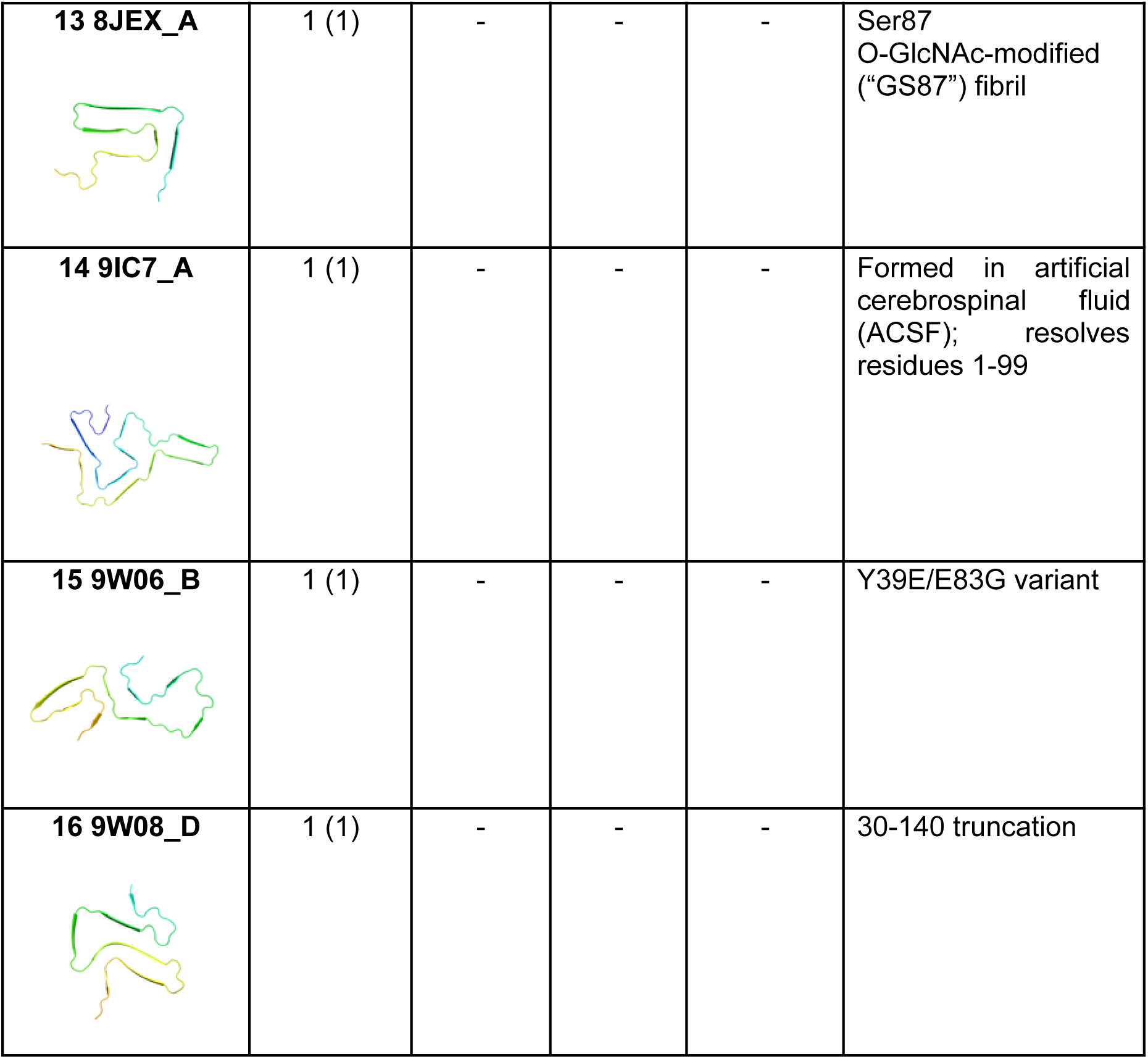
α-Synuclein polymorphs defined by TM-score clustering of the PDB. The 224 representative chains were clustered by average linkage at a TM-score distance of 0.5. For each cluster, the table gives the number of chains (and structures), the medoid (the chain with the highest mean TM-score to all other members), the median TM-score of members to the medoid, the most distant member and its TM-score to the medoid, and features shared by members. Singletons are chains whose average linkage to every cluster falls below a TM-score of 0.5, and are numbered 9-16.

| Cluster | No. chains (structures) | Medoid | Median TM-score to medoid | Most distant (TM-score) | Features |
| --- | --- | --- | --- | --- | --- |
| **Cluster 1** | 112 (99) | 9JD7_A | 0.70 | 9CDA_K (0.57) | Mainly pH 7.4; includes 6CU7 (pH 3) and juvenile-onset synucleinopathy-extracted 8BQW |
| **Cluster 2** | 67 (62) | 7OZH_A | 0.74 | 8CYY_I (0.58) | Highest proportion pH ≤ 6.5; includes E46K 6UFR and lipid-bound 8A4L, 8ADS, 8AEX |
| **Cluster 3** | 12 (12) | 8X7O_A | $\geq 0.97$ | 7C1D (0.70) | E46K (10/12); G51D 7E0F; E46K-seeded wild type |
| **Cluster 4** | 10 (1) | 2N0A_D | 0.54 | 2N0A_C (0.51) | Solid-state NMR structure 2N0A. "Most distant" here is the most divergent of its own 10 chains from the medoid chain, not a different structure. |
| **Cluster 5** | 5 (5) | 8ZLP_A | 0.77 | 6L1T_A (0.56) | N-acetylated polymorph 5A; pTyr39 |
| **Cluster 6** | 4 (4) | 9O9K_A | 0.90 | 8FPT<br>(0.54) | Lewy fold;<br>PD/Dementia with<br>Lewy Bodies (DLB)<br>patient-derived |
| **Cluster 7** | 4 (4) | 8ADU_A | 0.91 | 8JEY<br>(0.52) | Lipid-bound; pSer87 |
| **Cluster 8** | 2 (2) | 8GF7_A | 0.87 | 9D5C<br>(0.87) | O-GlcNAc Ser87 |
| **Singletons (8 chains, 7 structures - no cluster partner at TM ≥ 0.5)** |  |  |  |  |  |
| **9 6CU8_A** | 1 (1) | - | - | - | “Twister” polymorph<br>(Li *et al.*, 2018a),<br>full-length<br>recombinant, pH 3 |
| **10 7LC9_A** | 1 (1) | - | - | - | N-terminally<br>truncated (41-140)<br>construct |
| **11 7LC9_F** | 1 (1) | - | - | - | Same structure as<br>7LC9_A; a second,<br>distinct chain<br>conformation |
| **12 7V4C_A** | 1 (1) | - | - | - | Heparin-induced<br>polymorph 4 |
| **13 8JEX_A** | 1 (1) | - | - | - | Ser87<br>O-GlcNAc-modified<br>("GS87") fibril |
| **14 9IC7_A** | 1 (1) | - | - | - | Formed in artificial<br>cerebrospinal fluid<br>(ACSF); resolves<br>residues 1-99 |
| **15 9W06_B** | 1 (1) | - | - | - | Y39E/E83G variant |
| **16 9W08_D** | 1 (1) | - | - | - | 30-140 truncation |

The dominant structural clusters of α-synuclein are cluster 1 and 2. In fact, 179/224 chains fit into these definitions alone. Principal component analysis (PCA) demonstrates this, where the two major clusters separate clearly along PC1 (accounting for 64.3% of the variance, Figure 4). The remaining clusters sit between clusters 1 and 2 along PC1 but also separate from both dominant groups along PC2 (16.9% of the variance) (Figure 4). Overlaying properties including pH and protein source reveal some cluster differences, but no individual property explains the distribution of structures.

**Figure 4.**
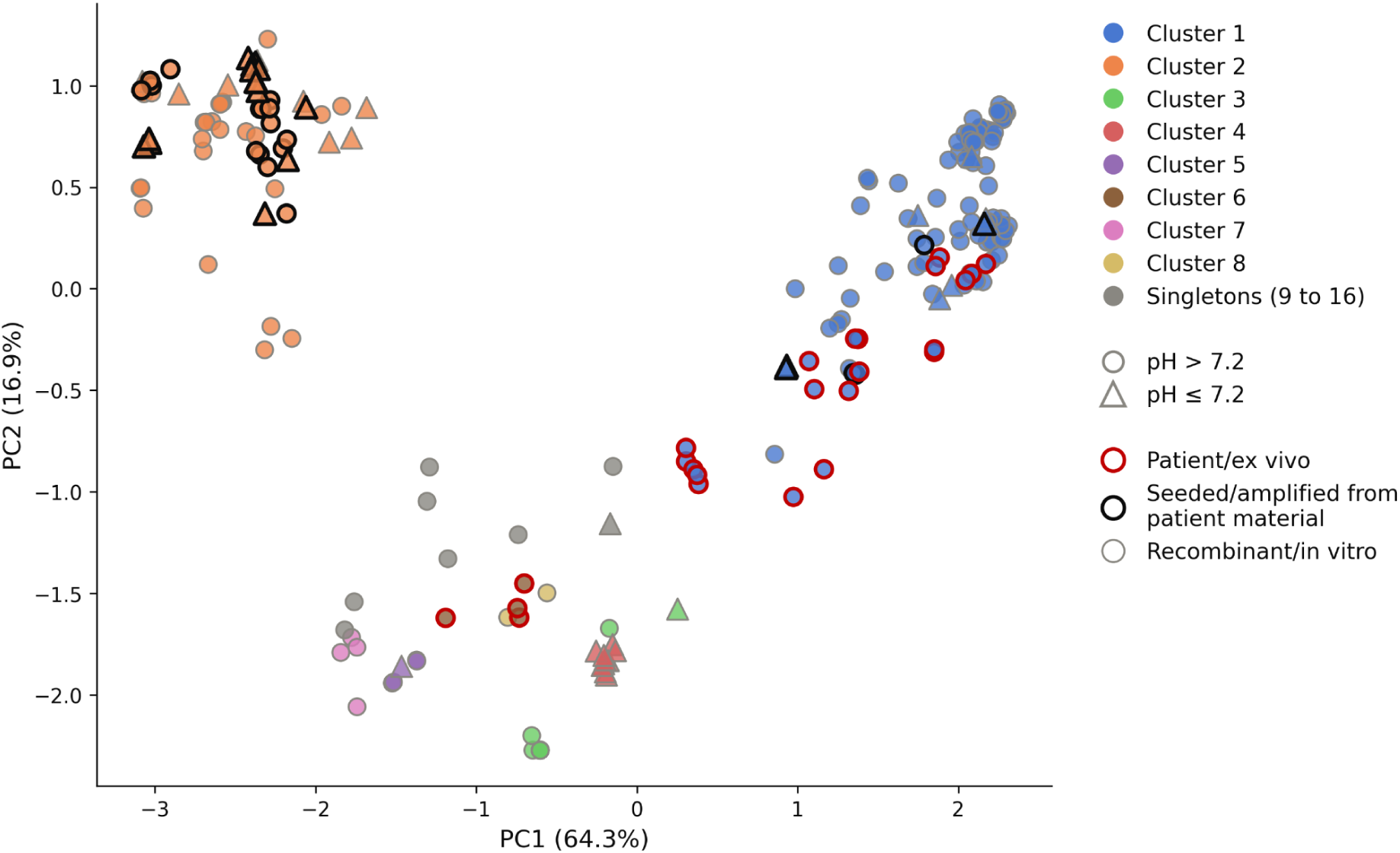
Principal component analysis of α-synuclein fibril structures in the PDB. PCA of the TM-scores; each point is one chain. Fill colour shows cluster, with singletons in grey. Shape shows fibril-formation pH: circles > 7.2, triangles ≤ 7.2. Outline shows source: red, patient-extracted; black, patient-seeded; grey, recombinant.

Cluster 1 is the largest, containing 112 chains, and exhibits the greatest internal structural variance, with a median TM-score to the medoid of 0.70 (Table 2, Supplementary Figure 1A). Despite this, the overall morphology of the polymorph group is similar, and the primary differences are localised to the termini. The core structure, spanning residues 43 to 81, is highly similar. In the member most distant from the medoid, 9CDA_K (TM 0.57), more of the N-terminus is resolved than in other cluster 1 structures, and the C-terminal region adopts a slightly different conformation, increasing the distance between Glu46 and Lys80 to 6.2 Å compared to an average of 3.8 Å across the whole cluster (Supplementary Figure 1C).

Cluster 2 is the second largest, containing 67 chains from 62 structures, with a median TM-score to the medoid (7OZH_A) of 0.74 (Table 2, Supplementary Figure 2). Of the 62 structures, 18 were formed at pH 6.5 or lower, more than seen in any other cluster and fitting with previous literature that suggests this polymorph is associated with low pH (Frey et al., 2024), although physiological pH examples are still present. Cluster 2 also includes 6UFR, a recombinant E46K fibril that is sometimes classified as a separate polymorph because of differences in protofilament stacking. Because we compare the folds of single chains, 6UFR falls within cluster 2 (TM 0.74 to the medoid); the features that set it apart are largely quaternary (Frey et al., 2024), whereas here we focus on tertiary fold.

The smaller clusters are largely defined by sequence variants, post-translational modifications, cofactors or tissue source (Table 2, Supplementary Figure 3-9), although these properties do not always map uniformly to a single cluster. Cluster 3 adopts an extended fold distinct from the more compact folds of clusters 1 and 2. The majority of the structures are of the hereditary E46K mutant, which accounts for 10 of the 12 members.The remaining two are 7E0F, a G51D fibril, and 7C1D, wild-type α-synuclein cross-seeded *in vitro* with E46K fibrils, which adopted an E46K-like fold (Long et al., 2021). Not all E46K structures fall in this cluster, however, as two further E46K fibrils, 6UFR and 8PJO, fall within cluster 2.

Modifications and cofactors map onto clusters, but less consistently than the sequence variants. Lipid-bound fibrils form their own cluster (cluster 7, Table 2), but three further lipid-bound structures from the same study sit within cluster 2, demonstrating lipids can promote multiple polymorphs (Frieg *et al*., 2022). Modifications at Ser87 are spread across cluster 7 (pSer87), cluster 8 (O-GlcNAc) and the singleton 8JEX. Cluster 5 brings together N-terminally acetylated and pTyr39 fibrils, while cluster 6, the Lewy fold, is the only group defined by tissue source, with all four structures derived from PD or DLB patients (Supplementary Figure 5-8).

Uniquely, cluster 4 is composed of 10 chains from a single solid-state NMR structure, 2N0A, that closely resembles that of cluster 1 (Table 2, Supplementary Figure 4). The eight singletons are similarly varied and are mostly recombinant, truncated constructs (Table 2, Supplementary Figure 9).

Clustering the α-synuclein PDB by TM-score recovers distinctions the field has already described (Scheres, Ryskeldi-Falcon and Goedert, 2023; Frey *et al*., 2024; Connor, Radford and Brockwell, 2025; Scheres, 2026), but gives them a quantifiable footing. It agrees closely with a recent objective classification (Milchberg *et al*., 2025), who found that 84% of around 100 structures fell into two classes, compared with 80% (179/224) in clusters 1 and 2 here. Our approach differs in two ways. Clusters are defined by TM-score’s established fold-equivalence threshold of 0.5, rather than by dataset-specific clustering parameters. Comparison is purely structural and down-weights variable termini, so constructs with different resolved ranges can be compared directly. The same criterion can also be applied directly to predicted models, which we use below to assign them to known polymorphs.

#### ABCFold for α-synuclein structure prediction

Having defined eight structural clusters and eight singleton α-synuclein polymorphs, we then asked which of these polymorphs current structure prediction methods can reproduce. To reduce chain clashes, we modelled only the fibril-forming core of α-synuclein (residues 37-97), as a monomer or oligomers of 1-15 chains, with AF3, Boltz-2 and OF3, run through ABCFold (v1.1.6), generating five models per method for each oligomer size, both with and without default template search. Models were excluded if they contained backbone clashes either within or between chains. As with the PDB structures, a single model can contain chains in different conformations, so chains within each model that differed by a Cα RMSD of at least 1.0 Å were separately retained to capture all polymorphs modelled. Each model was then compared against the 224 representative PDB chains with Foldseek in TM-align mode, and a chain was assigned to the cluster of its closest match if the TM-score was 0.5 or higher, and otherwise called novel.

For all methods, backbone clashes limited the number of usable models, and this varied depending on program, templates and oligomer size (Figure 5). This is not unexpected, as pathological amyloid arises from misfolding and differs markedly from the globular structures that dominate the methods’ training data. In terms of models without clashes, AF3 clearly outperforms the other methods, with 97% of models usable when templates are included, compared to 68% for the next highest method, Boltz-2. Without templates, AF3’s usable outputs drop to 68%, with Boltz-2 also achieving 68%, while OF3 produces only 67% and 43% usable models with and without templates respectively. Backbone clashes become more frequent as oligomer size increases. AF3 with templates produced fully usable models up to 12 chains, whereas without templates clashes appeared from 5 chains upward. Boltz-2 showed clashes from 4 chains onward regardless of templates, and OF3 without templates yielded no usable models at several sizes (7, 8, 9 and 12 chains).

**Figure 5.**
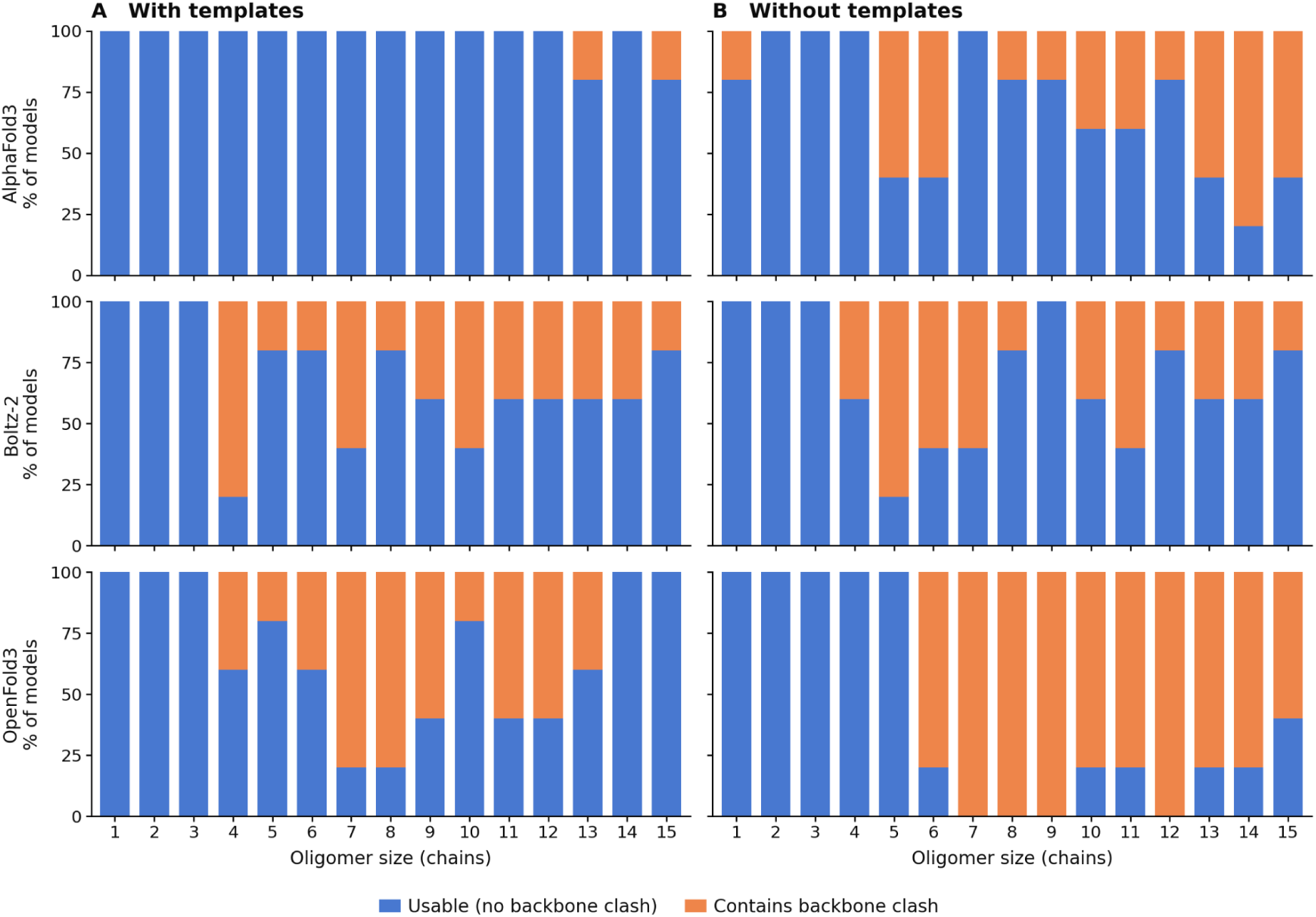
Backbone clash rate per oligomer size. Percentage of models that are usable (no backbone clash, blue) or contain a backbone clash (orange) for AF3, Boltz-2 and OF3, A) with and B) without templates. Five models were generated per method for each oligomer size.

Fibril-like structure was assessed for each model from two measures. The in-register stacking was quantified as the fraction of residues whose Cα was within 4.4-5.5 Å from the Cα of the equivalent residue in the neighbouring chain. This corresponds to a range around the 4.8 Å inter-strand spacing characteristic along a fibril axis. β-sheet content was assigned with DSSP (Kabsch and Sander, 1983). Models were called fibril-like if they contained at least two chains, 80% or higher stacked residues and 25% or more β-sheet content, with thresholds set by quantifying PDB structures. Whether a model was fibril-like depends heavily on the number of chains and the presence of templates, and pLDDT generally falls as the number of chains increases (Figure 6). With templates, AF3 produced fibril-like models uniformly from three chains upward. Boltz-2 and OF3 became fibril-like from five and four chains respectively, although OF3 was less consistent, returning non-fibril models at some sizes. Without templates, AF3 produced fibril-like models only between 7 and 11 chains and reverted to non-fibril structures at larger sizes, whereas Boltz-2 remained fibril-like from 5 chains upward irrespective of templates. Few OF3 models survived clash screening without templates, but all those at six or more chains were fibril-like.

**Figure 6.**
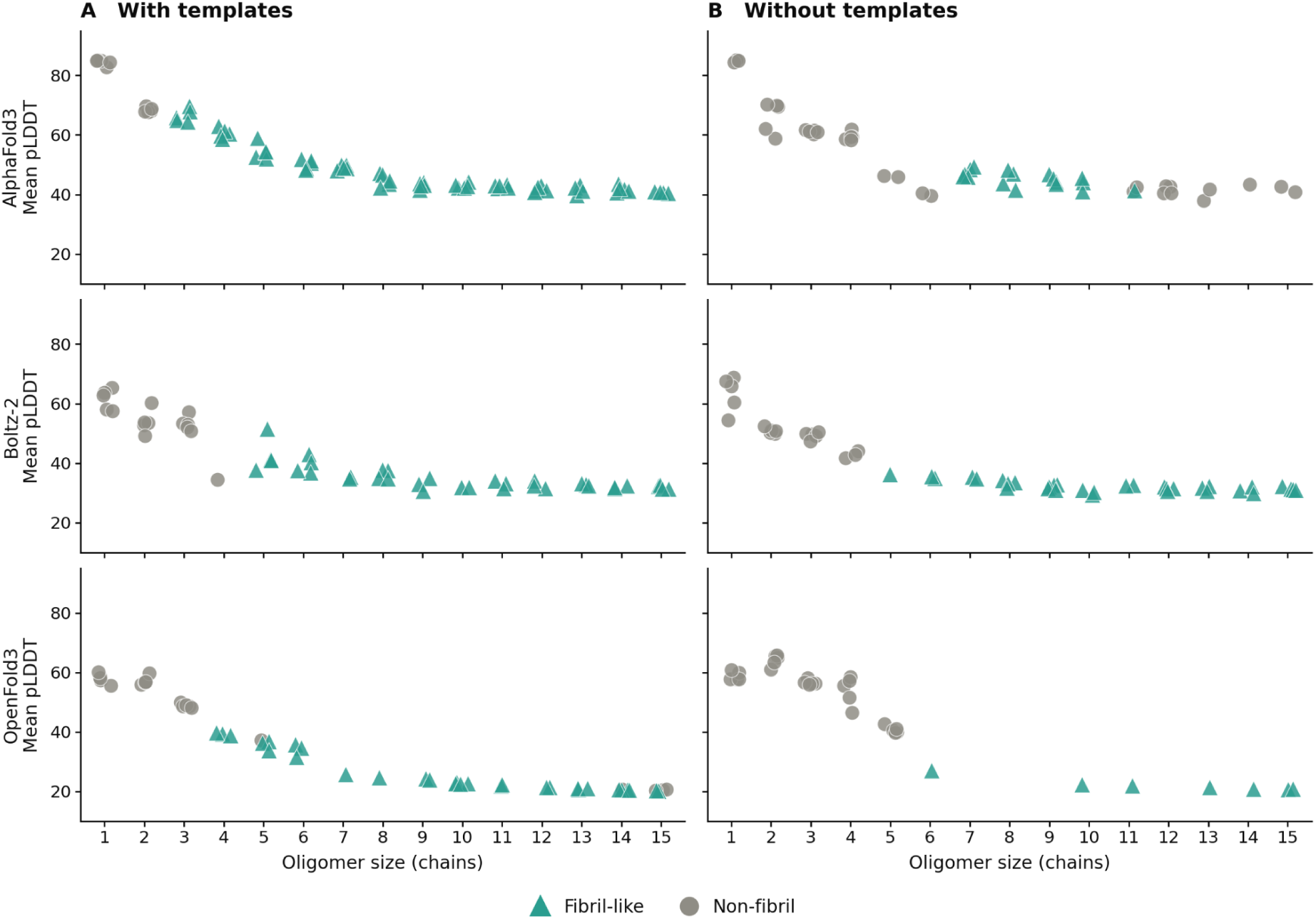
Mean pLDDT and fibril-like classification per oligomer size. Mean pLDDT of clash-free models for AF3, Boltz-2 and OF3, A) with and B) without templates. Fibril-like models are shown as triangles and non-fibril models as circles.

Of the eight clusters and eight singleton structures in the PDB, only six are represented as the closest match to a model, four clusters (1, 2, 6 and 8) and two singletons (11, 7LC9_F, and 16, 9W08_D) (Figure 7). This is based on each model’s closest match, but some models match more than one cluster or singleton with a TM-score above 0.5, particularly for clusters 1, 3 and 4, which are the most similar to each other. Fibril-like models also reproduced the handedness of experimental fibrils, 91% of which are left-handed. Many models were close to planar, with a twist below 0.5° (75% of AF3, 44% of Boltz-2 and 36% of OF3 models), but those with measurable twist were mostly left-handed (95%, 100% and 83%, respectively). Overall, models twisted about half as much as experimental structures (median model twist size 0.46° versus 0.93° for experimental).

**Figure 7.**
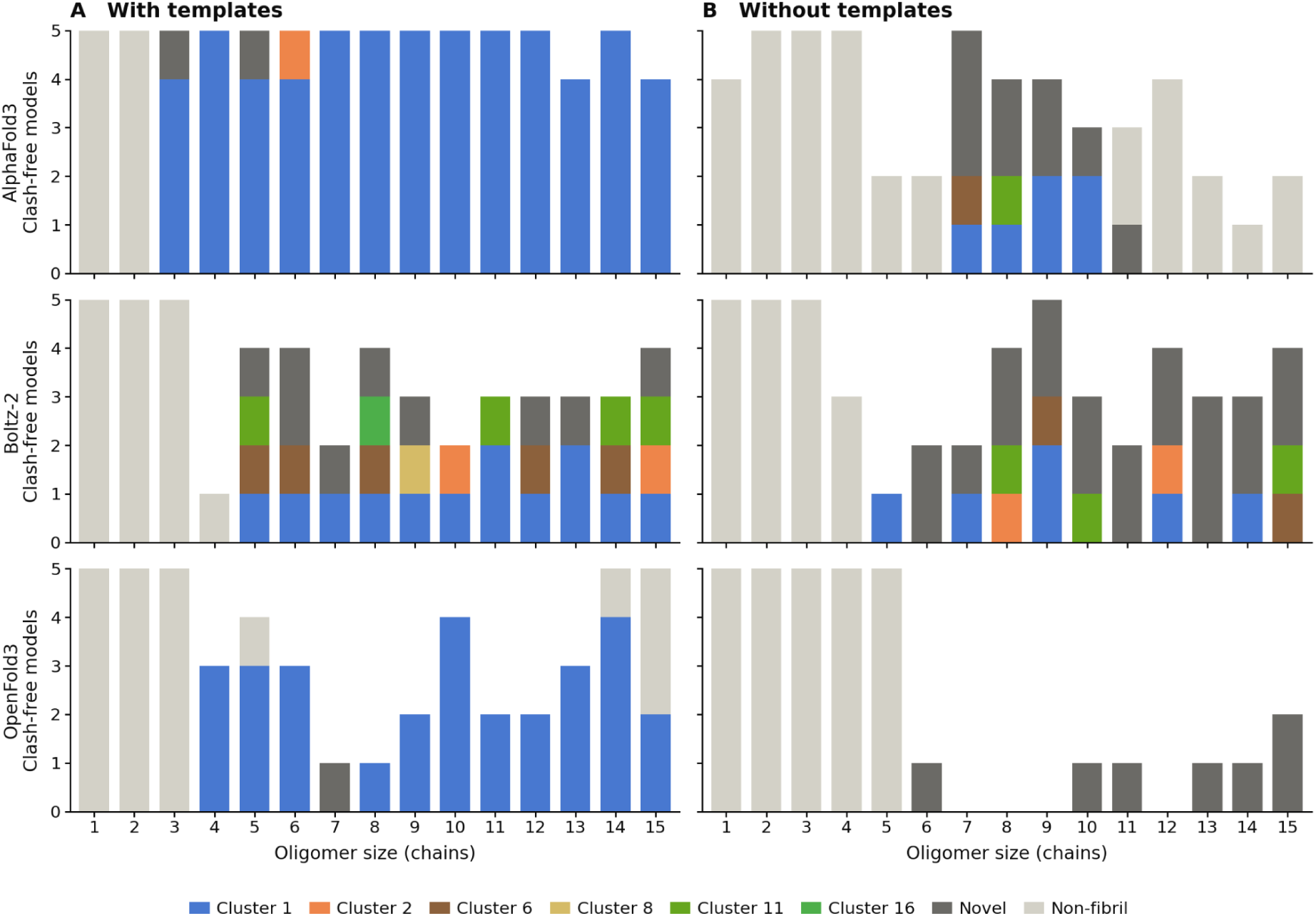
Closest PDB cluster match for each predicted model. Usable models from AF3, Boltz-2 and OF3, A) with and B) without templates, assigned to the PDB cluster or singleton of their closest match (TM-score ≥ 0.5), or classified as novel (fibril-like with no match at TM-score ≥ 0.5) or non-fibril. Models are shown by oligomer size and coloured according to cluster.

With templates, AF3 matched cluster 1 in 82% of usable models (60/73) and OF3 in 58% (29/50), with almost all remaining models from both methods classed as non-fibril or novel. Boltz-2 gave a broader spread, matching six different clusters and singletons across 26 of 51 models (51%), although 13 of these matched cluster 1. Without templates, cluster matches largely disappeared, with 8/51 AF3 models (16%) and 13/51 Boltz-2 models (25%) matching a cluster, compared to none of the 32 OF3 models. Most fibril-like models without templates were classed as novel. This includes 9/17 for AF3 (53%), 20/33 for Boltz-2 (61%) and all 7 OF3 models. These novel folds were confined to 6-15 chain oligomers (Figure 7B). With templates, AF3 and OF3 produced almost exclusively cluster 1 models. In contrast, Boltz-2 was less affected by templates and matched the widest range of clusters.

Several models achieve high TM-scores to the PDB, with the top matching model for each cluster shown in Figure 8. The single closest match was an OF3 15-chain model that aligned to 9JDK chain B with a TM-score of 0.91. High TM-scores are largely confined to cluster 1, however, with the highest match to any other group being a TM of 0.64 to the singleton 9W08 and to 7E0F in cluster 3. As our reference set includes structures deposited up to 2026, we could also ask whether any models resemble structures released after each method’s training cut-off. The singleton 9W08_D, as noted above, is matched at a TM-score of 0.64 by an OF3 4-chain model. 9W08_D is a truncated (30 to 140) fibril and a singleton here. The closest pre-cut-off structures, the cluster 1 members 6CU7_A and 8CE7_A, reach TM-scores of only 0.52 to 0.55 and share little residue-level correspondence with it (Cα RMSD above 10 Å, compared with 3.5 Å for the model; Figure 9A-B). A second example is an OF3 14-chain model that matches 9YFC_A, an A30P fibril deposited in September 2025, at a TM-score of 0.80, compared to 0.67 for the closest pre-cut-off structure, 6XYQ_B. 9YFC_A differs from earlier cluster 1 structures mainly at a kink at residues 59 to 60 and in the segment 73 to 79, and in both regions the OF3 model follows 9YFC_A (median Cα deviation 1.1 Å over residues 73 to 79, compared with 3.3 Å from 6XYQ_B, Figure 9C-D).

**Figure 8.**
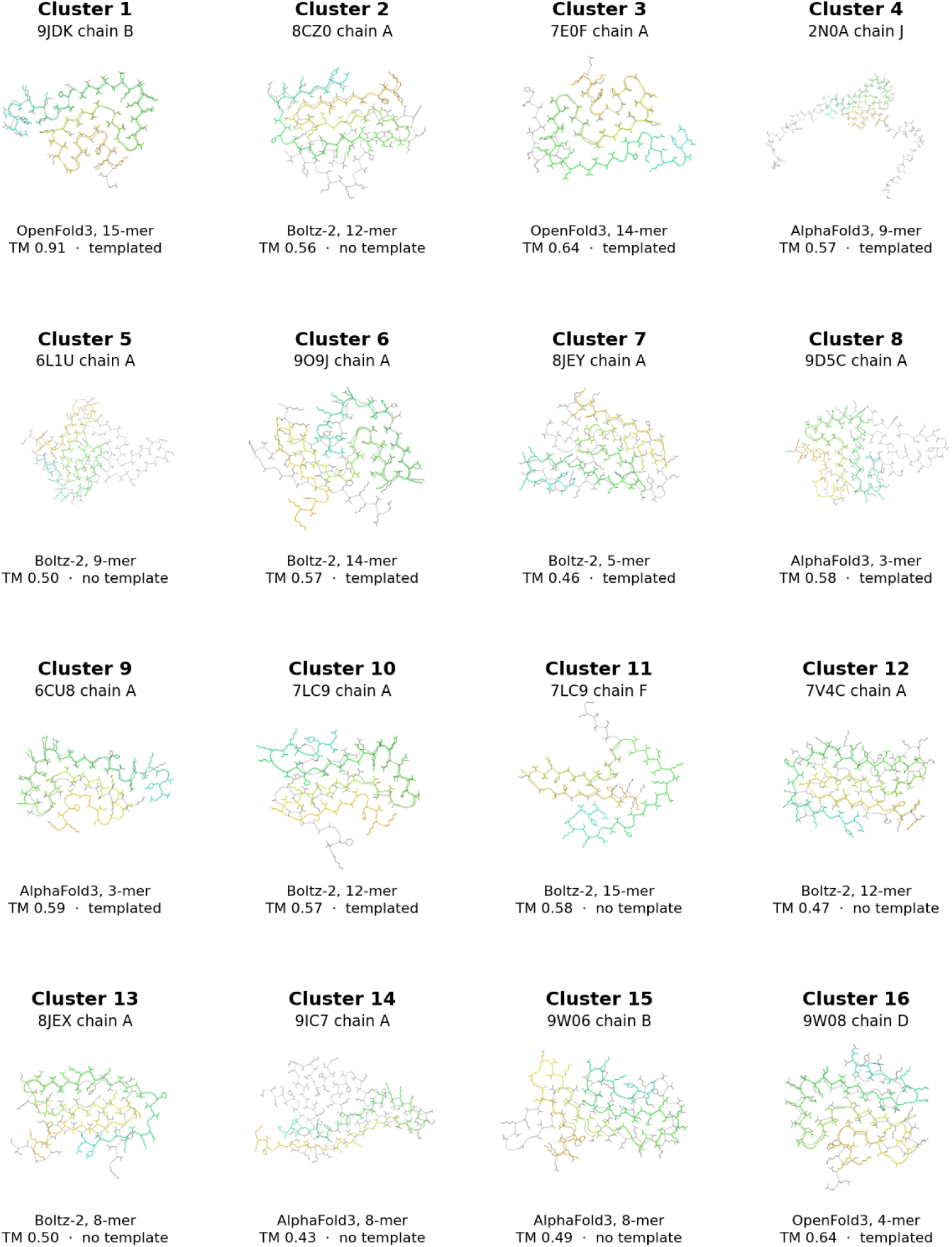
Best-matching predicted models for each α-synuclein PDB cluster. The highest-scoring model (by TM-score) for each cluster or singleton, superposed on its closest PDB chain. The prediction method, number of chains and TM-score are given for each pair. PDB structures are shown in grey, with models coloured rainbow from N to C (blue to red).

**Figure 9.**
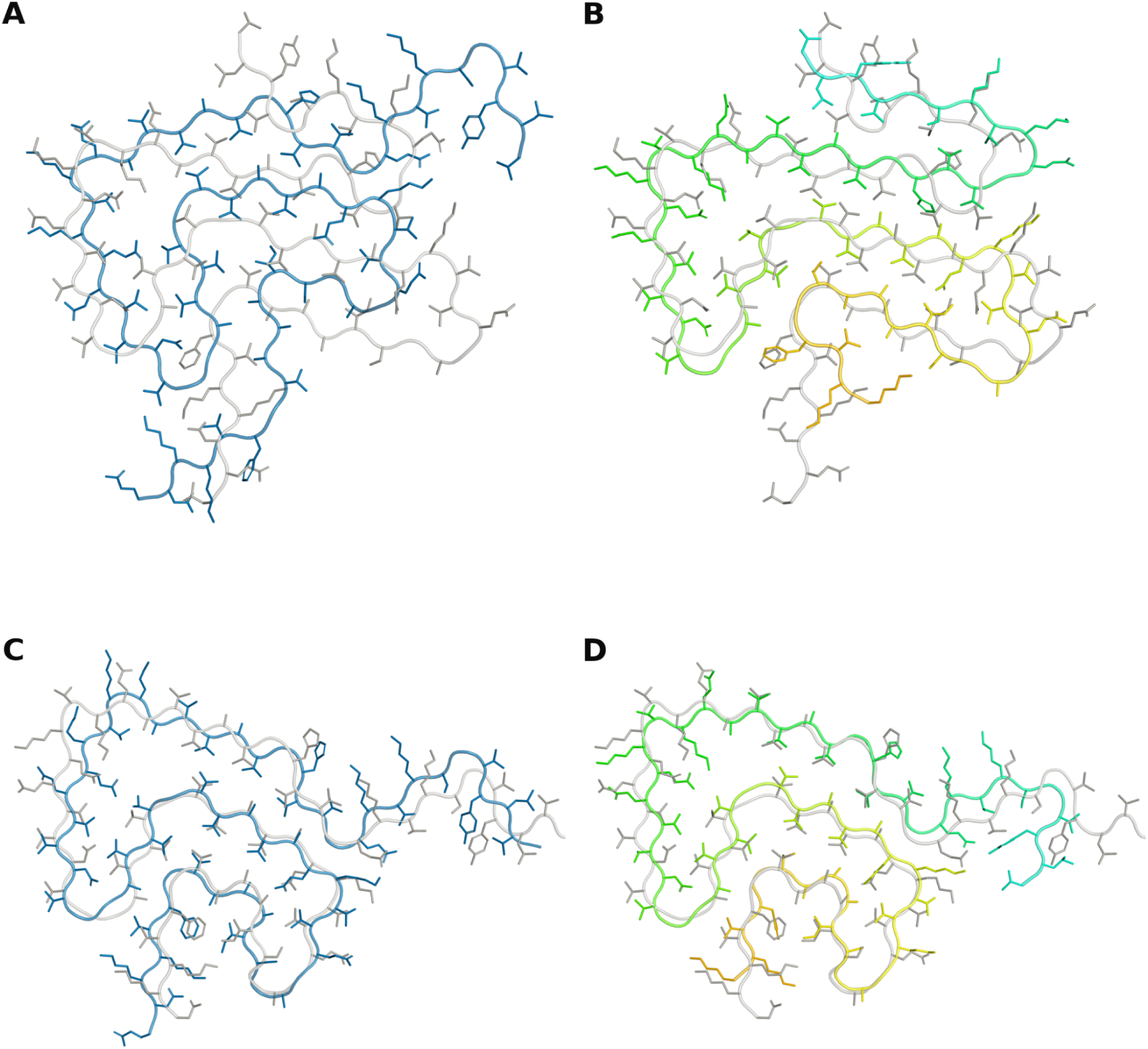
Predicted models resembling structures released after the training cut-off. Single chains viewed down the fibril axis, with the newer PDB structure in grey. A) 9W08_D with the closest earlier structure, 8CE7_A (blue; TM-score 0.55). B) 9W08_D with the closest OF3 model (rainbow; TM-score 0.64). C) 9YFC_A with the closest earlier structure, 6XYQ_B (blue; TM-score 0.67). D) 9YFC_A with the closest OF3 model (rainbow; TM-score 0.80).

Both post-cut-off structures differ from the closest earlier structure, and in both cases the OF3 models were substantially closer to the new structure (0.64 and 0.80) than any earlier structure (0.55 and 0.67). These methods can therefore produce models that resemble newly determined polymorphs more closely than any structure they were trained on. For 9YFC_A, a cluster 1 member, the model appears to recombine variation already present in the cluster, matching the kink at residues 59 to 60 and the conformation of residues 73 to 79 (Figure 9D).

Several of the models produced are fibril-like but do not match any known α-synuclein structure. Clustering these by TM-score, in the same way as the PDB structures, separates the 82 novel chains into 12 clusters and six singletons. Novel folds were found across a range of oligomer sizes, both with and without templates, and five of the 12 clusters contained models from more than one method. However, models were generally more similar to each other within a single method than between methods. TM-scores between models of the same method reached as high as 0.88, whereas the best TM-score between models of different methods reached only 0.70 (AF3/Boltz-2) (Figure 10A), achieved independently by two separate method pairs. Although there is some agreement between methods, it is weaker and less common than within a single method, so is unlikely to represent one well-defined alternative fold. Some of these models are also implausible for full-length α-synuclein, as the N- and C-termini are buried within the fold.

**Figure 10.**
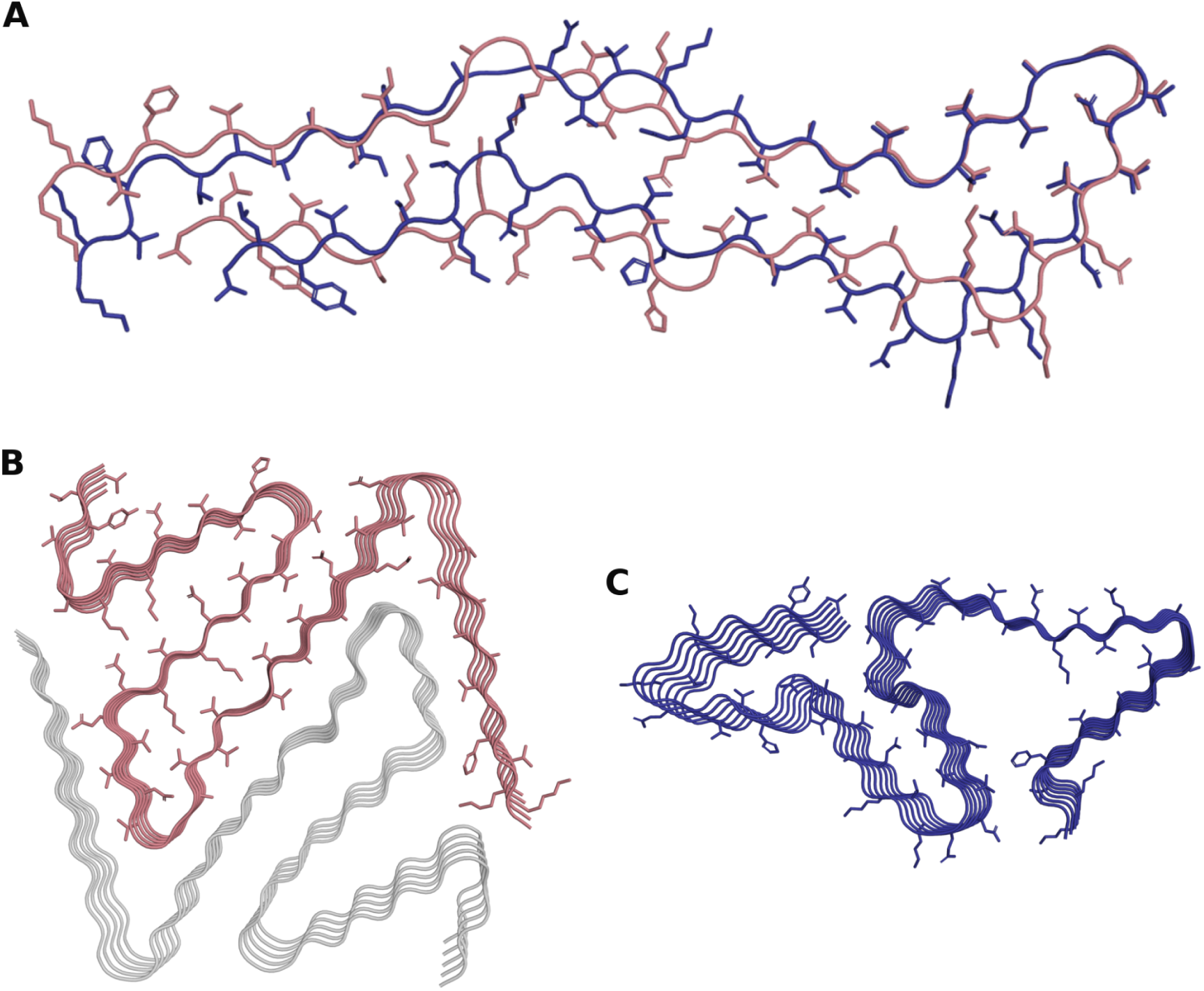
Novel fibril-like folds predicted for α-synuclein. A) The closest pair of novel models from different methods: AF3 (blue, 8 chains) superposed on Boltz-2 (salmon, 15 chains), TM-score 0.70. B, C) Novel models whose rise and pitch fall within the range of solved α-synuclein fibrils, viewed down the fibril axis. B) Boltz-2, 10 chains (rise 4.75 Å, pitch 177 nm), forming two protofilaments (salmon and grey). C) AF3, 7 chains (rise 4.87 Å, pitch 203 nm), forming a single protofilament. All models were predicted without templates.

We next compared the helical geometry of the novel models with that of solved α-synuclein fibrils (pitch 110-280 nm, rise ∼4.8 Å). Rise is constrained by cross-β hydrogen bonding, and most novel models (87%) fell within this range. Pitch is less constrained, because the observed range partly reflects the dependence of cryo-EM on twist, and most models had longer pitches than solved fibrils. 15 of the 46 novel models matched solved fibrils on both measures, including examples from more than one method. These models therefore resemble known α-synuclein fibrils in their stacking geometry and may represent polymorphs not yet determined experimentally. The model in Figure 10B forms two protofilaments, so the large cavity apparent in a single chain is occupied by the second protofilament. The model in Figure 10C encloses an internal cavity within a single protofilament. Cavities also occur in experimental α-synuclein fibrils, including a hydrophilic tunnel containing unassigned density within the protofilament core (Guerrero-Ferreira *et al*., 2018) and cavities enclosing non-proteinaceous molecules in filaments from multiple system atrophy brain (Schweighauser *et al*., 2020).

#### Capturing α-synuclein polymorphism with deep-learning based structure prediction

Polymorphs differ in the surfaces they expose, so interactions with small molecules, antibodies and other proteins can be polymorph-specific, and ligand design needs to account for which polymorphs are present (Li *et al*., 2018b; Milchberg *et al*., 2025). Therefore, the ability to model a specific polymorph is required across many applications. Modelling ligand binding, post-translational modifications or mutations in the context of a particular fold therefore depends on reproducing that fold reliably.

To test whether each method could recreate all known α-synuclein polymorphs, the fibril-forming core was modelled with each of the 16 reference structures (eight cluster medoids and eight singletons) supplied as a template. Templates were combined with MSA depths of 1, 2, 4, 8, 16, 32 and the full MSA (71 sequences by MMseqs2). Oligomers of 1 to 8 chains were modelled. In the earlier runs, templated models from all three methods were fibril-like by six chains, so this range spans the transition from single chains to fibril-like stacks for all methods. For every combination of target polymorph, method, oligomer size and MSA depth, each predicted chain was scored against the template it had been given, and the highest TM-score across the five models of that condition was taken. A fold was counted as recovered when this value reached the TM-score threshold of 0.5 used throughout.

When pooled across all conditions, AF3 and Boltz-2 each recovered 13 of the 16 target folds, whereas OF3 recovered only three and never exceeded a TM-score of 0.57 to any template (Figure 11A, Supplementary Figure 11). AF3 did not recover clusters 3, 9 and 11, and Boltz-2 did not recover clusters 9, 11 and 14, leaving singletons 9 and 11 as the only targets that every method failed to produce. These are the twister polymorph 6CU8_A (Li *et al*., 2018) and 7LC9_F, one of two distinct chain conformations in an N-terminally truncated fibril. Both templates cover only part of the modelled core (residues 43 to 83 for 6CU8_A and 61 to 98 for 7LC9_F), and both folds depend on contacts that a single templated chain cannot provide: 7LC9_F packs against the differently folded chains of the same fibril through an asymmetric interface, while 6CU8_A differs from the rod polymorph of cluster 1 in its protofilament interface (Li *et al*., 2018).

**Figure 11.**
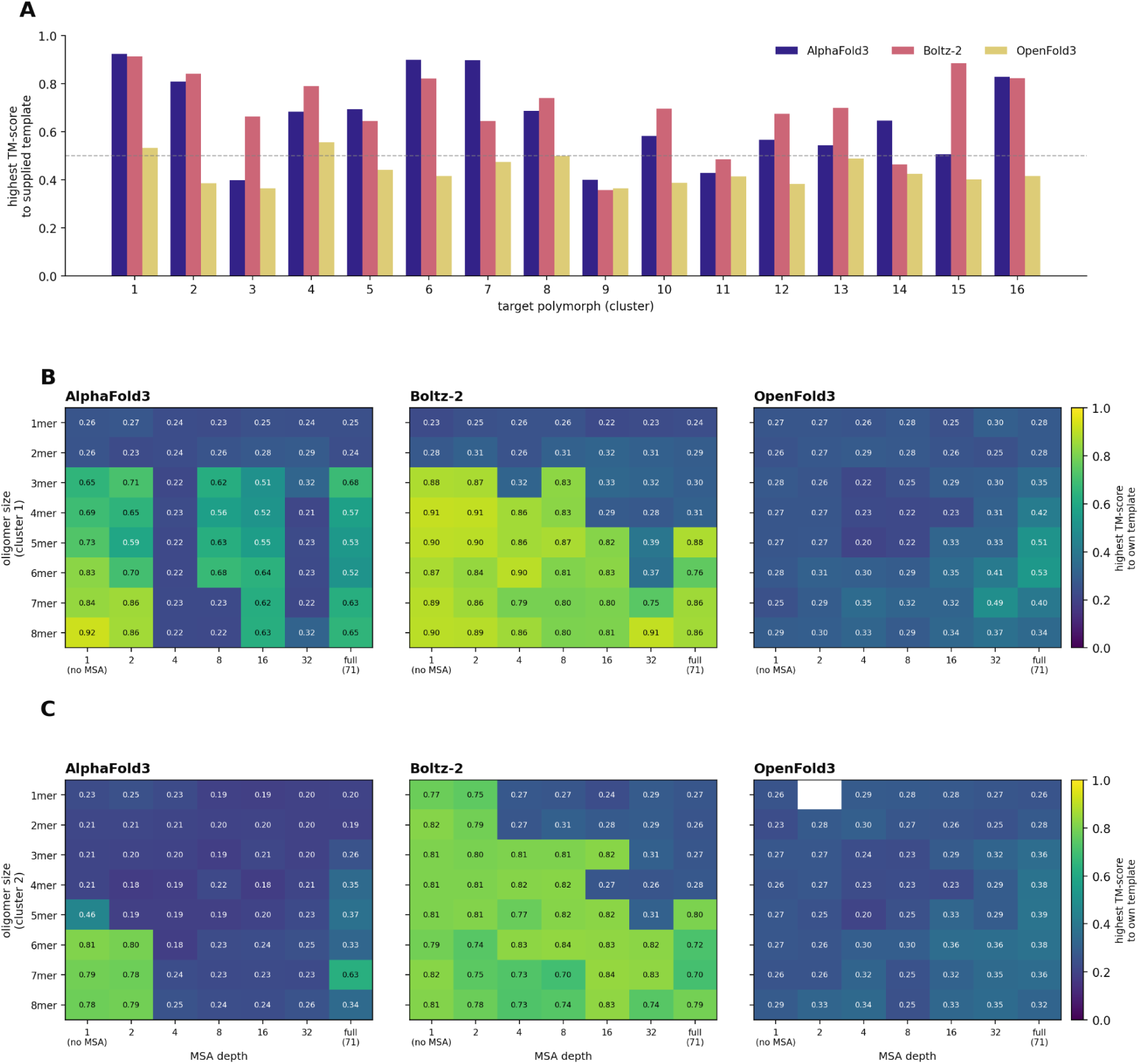
Template obedience for each α-synuclein polymorph. A) Highest TM-score to the supplied template for each of the 16 target folds, across oligomer sizes and MSA depths, for AF3, Boltz-2 and OF3. The dashed line marks the TM-score 0.5 fold-equivalence threshold. B) TM-score to the supplied template by oligomer size and MSA depth for cluster 1 and C) cluster 2. Five models were generated for each combination of method, oligomer size and MSA depth.

Although AF3 and Boltz-2 recover a similar number of folds, they do so under different conditions. Boltz-2 matches its template with fewer chains. For cluster 2, Boltz-2 reaches a TM-score of 0.77 as a single chain and changes little up to eight chains, whereas AF3 needs six chains to recreate the fold. For cluster 1, both methods reach the template at three chains (0.88 for Boltz-2, 0.71 for AF3). However, Boltz-2 then remains near 0.90, while AF3 continues to improve with oligomer size, reaching 0.92 at eight chains (Figure 11B). OF3 shows no comparable improvement for any target. This suggests AF3 relies more on chain numbers than Boltz-2 to follow a template.

The effects of chain number and MSA depth are also linked (Figure 11B-C, Supplementary Figure 10). For a given target and method, the template is only reached once a certain number of chains is modelled, and this number generally increases with MSA depth. Boltz-2 models of cluster 1 reach the template at three chains with a single sequence input, at four chains with a depth of four, at five chains with a depth of 16 and at seven chains with a depth of 32. Cluster 2 behaves the same way, reaching the template from a single chain at MSA depths of one and two, from three chains at depths of four to 16, and from six chains at a depth of 32. Additional chains therefore compensate for a deeper MSA, and a shallow MSA compensates for fewer chains.

AF3 shows the same trade-off across a narrower range. At a depth of four sequences, cluster 6 remains near 0.20 up to seven chains before reaching 0.73 at eight, and cluster 16 at a depth of 32 recovers only at seven chains. For some targets no number of chains compensates, with cluster 2 remaining near 0.20 at every size once the depth reaches four. OF3 shows no equivalent behaviour for any target at any combination. This fits with the amyloid fold not being the conformation encoded in the MSA by natural selection, meaning evolutionary information works against both the template and the stacking from additional chains. However, for several AF3 targets, including clusters 2, 4 and 9, TM-scores partly recover at full MSA depth after falling at intermediate depths.

## Discussion

The release of AF3 and related methods has opened new avenues for predicting protein complexes, including the particularly challenging case of amyloid assemblies. While AF2 reproduced monomeric α-synuclein structures, its multimeric predictions consistently failed to adopt a fibril architecture, and even with custom parameters a fibril fold was only recovered for a monomer. This is notable given the tendency of AF2 to produce confident but unrealistic β-sheet rich structures (Pratt *et al*., 2025). AF3, by contrast, produced fibril-like α-synuclein trimers using default settings.

Running AF3, Boltz-2 and OF3 through a common pipeline showed that they behave differently, and that no single method is best for every purpose. AF3 produced the fewest clashes and the most consistently fibril-like assemblies, but with templates it returned almost exclusively cluster 1 folds. Boltz-2 was less constrained by templates and matched the widest range of known polymorphs. OF3 performed worst overall and recovered only three of the 16 targets when templated, yet it produced models resembling structures released after the training cut-off. Sampling a conformational landscape as diverse as amyloid polymorphism therefore benefits from running several methods.

The template results give a practical route to modelling a chosen polymorph. Across methods, following a template depended on both the number of chains modelled and the depth of the MSA, with the two trading off against each other: additional chains compensated for a deeper alignment, and a shallow alignment compensated for fewer chains. This fits with the amyloid fold not being the conformation encoded in the MSA by natural selection, so that evolutionary information competes with both the template and the stacking imposed by neighbouring chains.

Amyloid polymorphs are defined as structural variants of amyloid fibrils formed by the same protein sequence but adopting different conformations (Sawaya *et al*., 2021; van der Kant *et al*., 2022; Connor, Radford and Brockwell, 2025). Polymorphism can be influenced by various factors including post-translational modifications, cofactors and co-aggregation and disordered regions outside of the fibril core (Li and Liu, 2023; Aubrey and Radford, 2025; Watts, 2025; Pradhan *et al*., 2026). Amyloid polymorphism is important as different polymorphs are associated with different disorders, as well as differences in disease progression (Sawaya *et al*., 2021; Gomez-Gutierrez *et al*., 2023). Despite their importance, there is no objective consensus on what level of difference is classed as a polymorph. Here we have shown that TM-score, although designed for globular proteins, sorts the available structures into clusters that broadly agree with expert classifications of α-synuclein polymorphs, and with the independent clustering of Milchberg *et al*. (2025), while providing an objective threshold that can be applied consistently to new structures and to predicted models.

AF3’s ability to predict amyloid structures has important implications. Firstly, the potential to model polymorphs by AF3, including seen and unseen polymorphs, allows exploration of structural diversity in amyloid proteins without solely relying on experimental methods or on computationally expensive molecular dynamics simulations, something that is established for globular proteins. While cryo-EM has significantly advanced amyloid structure determination it has inherent limitations (Li *et al*., 2013; He and Scheres, 2017b). Cryo-EM can resolve major polymorphs, but for species that are less abundant there may be too few particles available for reliable reconstruction (Lutter, Aubrey and Xue, 2021; Zielinski, Röder and Schröder, 2021). In a similar vein, obtaining sufficient quantities of patient-derived fibrils for reconstruction is often challenging (Sanchez *et al*., 2024). Finally, non-twisting fibrils cannot currently be resolved with helical reconstruction tools (Zielinski, Röder and Schröder, 2021; Sanchez et al., 2024). Therefore, it is common to only determine the structure of a fraction of polymorphs. It should also be noted, finally, that extraction methods often require homogenisation and solubilisation/separation methods that could be biasing accessible fibril structures or removing key associated cofactors. Thus, novel folds identified via modelling could represent currently unidentified folds that are intractable with current mechanisms of extraction and identification. AlphaFold models are already commonly used in globular structure resolution (Millán *et al*., 2023; Alshammari, He and Wriggers, 2025) but this has yet to be applied to fibril structures. We envisage there will be cases where helical reconstruction of filaments could be aided by AF3-generated candidate fibril models.

Twist was the feature least well reproduced by any method. Models recovered the handedness of experimental fibrils, but most were close to planar and twisted about half as much as solved structures. This is unlikely to be learned from the training data, which is dominated by twisted fibrils. Every cryo-EM structure in our reference set has a measurable twist. The only planar stacks in the PDB are solid-state NMR models, in which twist is not directly determined. The flatness of the models more likely reflects how current methods assemble short stacks of chains without imposed helical symmetry, and twist varied with oligomer size in a method-specific way, unlike in the PDB. Methods that model helical twist explicitly, whether by imposing helical symmetry or by modelling longer stacks, may both improve the accuracy of individual folds and broaden the range of polymorphs sampled.

The ability of AF3 to sample broad structural diversity is particularly relevant for α-synuclein, where patient-derived fibrils show diverse folds. Three types of folds have been identified in patient-derived samples from synucleinopathies corresponding to DLB, JOS and multiple system atrophy. Two types of fibril structures have been extracted from multiple system atrophy brains and have similarities to those extracted from JOS patients. Here, the multiple system atrophy and JOS folds fall within cluster 1, while the Lewy fold, extracted from DLB and PD brain, forms cluster 6 (Yang *et al*., 2023). Twist also contributes to structural diversity; Lewy-fold fibrils have a right-handed twist, whereas multiple system atrophy fold fibrils twist left-handedly (Schweighauser *et al*., 2020; Yang *et al*., 2022); Yang et al. (2022) explored comparing non-twisted fibrils with twisted fibrils using 2D class average comparisons of projections to identify similarities between folds. Having access to reliable fibril models could assist with this process and expand the feasibility of generating structural models of additional fibril morphologies.

Beyond disease-specific folds, mutations also shape the fold: most E46K structures form cluster 3, although two fall within cluster 2. New mutations for a range of disease-associated amyloid proteins are being identified, but for many the effect of the mutation is not clear. Modelling could provide an avenue to explore how mutations could lead to disease and how they could be targeted differently. This is important given that disease onset, symptoms and duration vary between different mutations, and so therapeutic targeting needs to consider genetic variants as isolated diseases. The ability of AF3 and similar programs to sense the impact of mutations is generally considered limited: mutant structures often strongly resemble wild-type (Buel and Walters, 2022), although local confidence can indicate that a new side chain is unfavourably placed (Pak *et al*., 2023). However, it is worth recalling that models of amyloid will not be driven by evolutionary covariance information in the same way: they may therefore be more sensitive to mutation, and we are exploring this hypothesis.

Our modelling has included the wild-type α-synuclein sequence in the absence of any post-translational modifications but it is known that α-synuclein can be modified in several ways including phosphorylation and nitration (He, Yu and Chen, 2019; Hu *et al*., 2024; Kochen *et al*., 2024). Elucidating the effect of modifications experimentally can be difficult due to requiring tightly controlled environments to achieve complete modification and/or single site modifications. Modelling could provide a platform to further understand these processes. Our clustering shows that the presence of modifications map onto specific folds: lipid-bound and pSer87 fibrils form cluster 7, O-GlcNAc Ser87 fibrils cluster 8, and N-terminally acetylated and pTyr39 fibrils cluster 5. Modelling the structural consequences of a modification therefore requires the relevant polymorph, which the template-guided results suggest is achievable for most known folds.

Cofactors further modulate fibril structure and have been identified in many α-synuclein folds. In vitro aggregation studies (Cohlberg *et al*., 2002; Ohgita *et al*., 2025) have explored the effect of cofactors on aggregation rate but there is still a lack of understanding how these cofactors affect pathological progression of disease. This is particularly evident in the apparent association between metal exposure and PD (Zhao *et al*., 2023). Understanding how cofactors e.g. metal ions are integrated within fibril structures could provide information relevant to disease progression and therapeutic targeting. AF3’s ability to include ligands offers a route to screening cofactor association systematically, and combining ligands with a polymorph-specific template would allow such screens to be carried out in the context of a defined fold rather than whichever conformation a method produces by default.

A key area of research not fully understood is how protein structure and cell death are related to the pathological spread of amyloid throughout the brain and other organs in systemic amyloid diseases. Given the lack of correlation of patient-derived fibril structures with those from experimental models it is currently difficult to map the aggregation process in vivo, and particularly the influence of cofactors and environmental conditions. Oligomeric or pre-fibrillar species are often considered to be the important feature in amyloid pathogenesis (Wells *et al*., 2021). However, their transient nature means they are often experimentally intractable. Establishing reliable modelling methods could provide a route to generating candidate structures for these species, for example by modelling small oligomers of the aggregation-prone region and asking which arrangements are predicted consistently, or by exploring how co-aggregation or cross-seeding with a second sequence alters the fold adopted. Our oligomer series shows that predictions change with the number of chains modelled, which may itself be informative about the point at which a stable cross-β arrangement becomes favourable.

These predictions have the translational application of aiding in the development of fibril targeting therapeutics. Accurate structural models of different polymorphs can inform the design of cross-polymorph inhibitors that target conserved structural motifs common to all fibrils, as well as polymorph-specific targeting of disease-relevant or patient-specific polymorphs. The same applies to diagnosis. Distinguishing synucleinopathies clinically is difficult, particularly early in disease. Seed amplification assays have been shown to discriminate PD from multiple system atrophy using cerebrospinal fluid, with the aggregates from each corresponding to different conformational strains (Shahnawaz *et al*., 2020). Imaging ligands that bind α-synuclein fibrils have also been developed but do not yet distinguish between polymorphs. Structural models of individual polymorphs could support the design of fold-selective tracers and the interpretation of such assays.

Further understanding of amyloid polymorphism may help to explain disease heterogeneity, guiding patient-driven design of therapies. It has previously been suggested that specific folds of neurodegenerative amyloid proteins define distinct diseases, whilst identifying that several conditions can also share the same fold (Scheres, Ryskeldi-Falcon and Goedert, 2023). It has also been proposed that different diseases could arise from different protofilament assembly of a common core fibril structure. Where multiple filament types have been identified in an individual they may also share a common fold, differing in protofilament assembly or be formed from distinct protofilament folds (Schweighauser *et al*., 2020). Where multiple protofilament folds exist, the proportion of the different folds could be related to disease severity and brain location. The ability to model interactions of specific folds could therefore develop pipelines for disease-specific modulation which is currently difficult to do using existing experimental model systems that often do not reproduce patient-derived fibril structures or assembly pathways. In comparison to tau where fibril structures exist for many of the different tauopathies, there are relatively few patient-derived α-synuclein fibril structures available (Schweighauser *et al*., 2020).

We have shown AF3 is able to generate feasible fibril models in a way not possible with earlier versions of AF. However, as shown in Figure 2B, AF3 does not model the pitch associated with fibril morphologies. This may limit the use of these models for structure determination and design, but it is also a tractable target: the open weights of Boltz-2 and OpenFold 3 make it possible to fine-tune on the growing set of amyloid structures, which may improve both the accuracy of individual models and the range of polymorphs sampled.

Model systems are helpful in understanding assembly processes and generating fibril structures e.g. from aggregation at non-physiological pH or cross-seeding wild-type proteins with mutant seeds. However, we note that inclusion of fibril structures that may be unlikely to exist within the body within large-scale modelling to identify novel polymorphs may cause issues. We therefore propose that future modelling approaches could include additional screening to remove fibril structures only observed in non-physiological conditions, and so enhance the translatability of modelling to pathological disease understanding.

Amyloid assemblies remain a new frontier for structure prediction. They appeared as targets in CASP17 for the first time, but the field has yet to establish a framework for working with this class of protein. The approach described here offers a starting point for amyloid structure research. Applied to α-synuclein, these measures show that current methods already sample real structural space. They can, with steering, reproduce most known α-synuclein polymorphs, generate models closer to newly determined structures than anything in their training data, and can produce plausible models that are beyond solved structures. This work provides practical guidance for modelling a fibril: running more than one method, pairing templates with varied alignment depths to reach a specific polymorph and varying the number of chains can all influence the structural outcome. Our findings support deep-learning structure prediction as a powerful tool for amyloid biology.

## Funding acknowledgements

BBSRC GIBA Fellowship to RP (UKRI775).

---

**Supplementary Figure 1.**
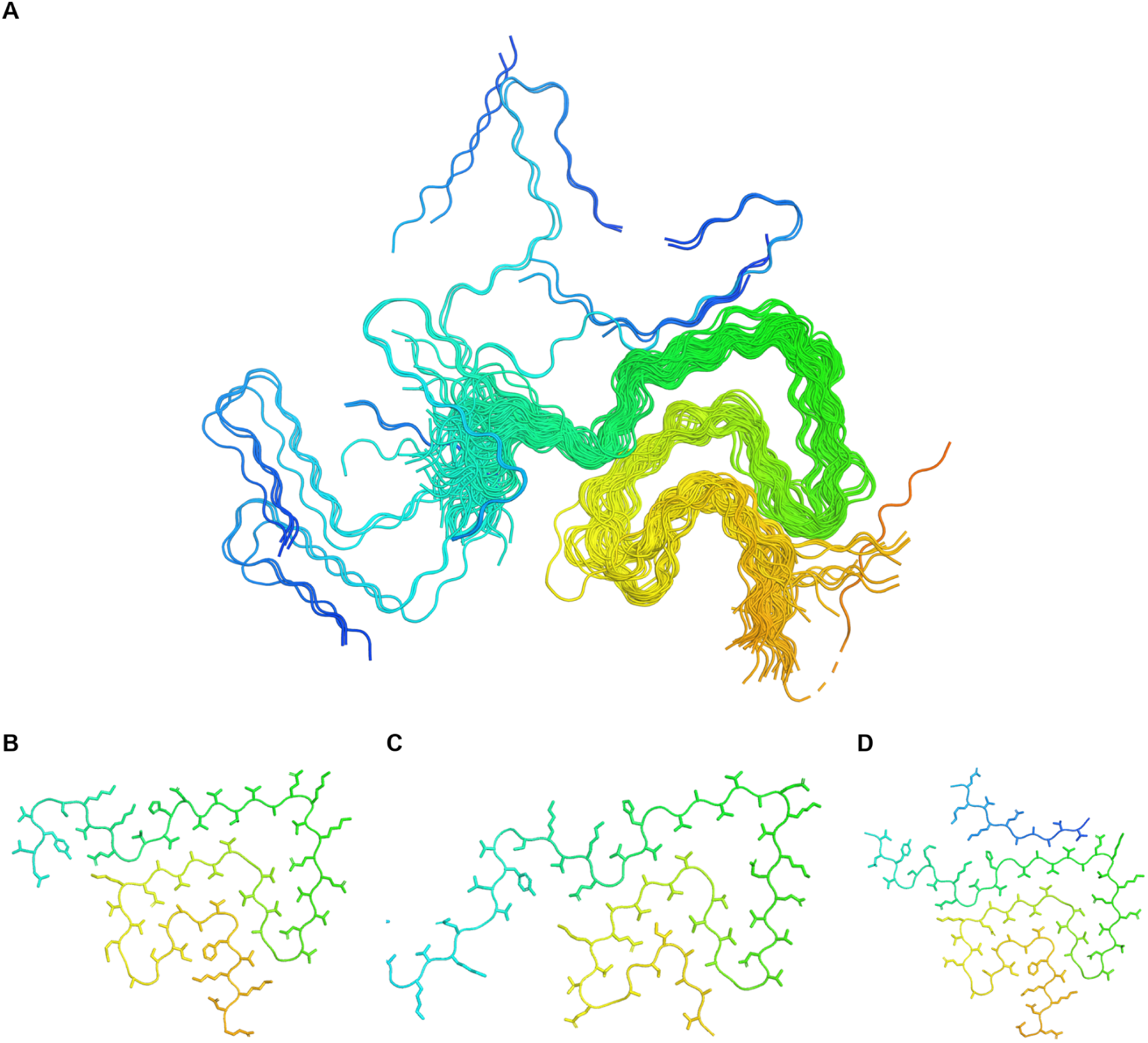
Cluster 1. A) Superposition of all cluster 1 chains, coloured N (blue) to C (red) terminus. B) Medoid 9JD7_A, C) the chain furthest from the medoid, 9CDA_K, and D) the JOS-extracted α-synuclein structure 8BQW_A, shown as single chains with side chains displayed.

**Supplementary Figure 2.**
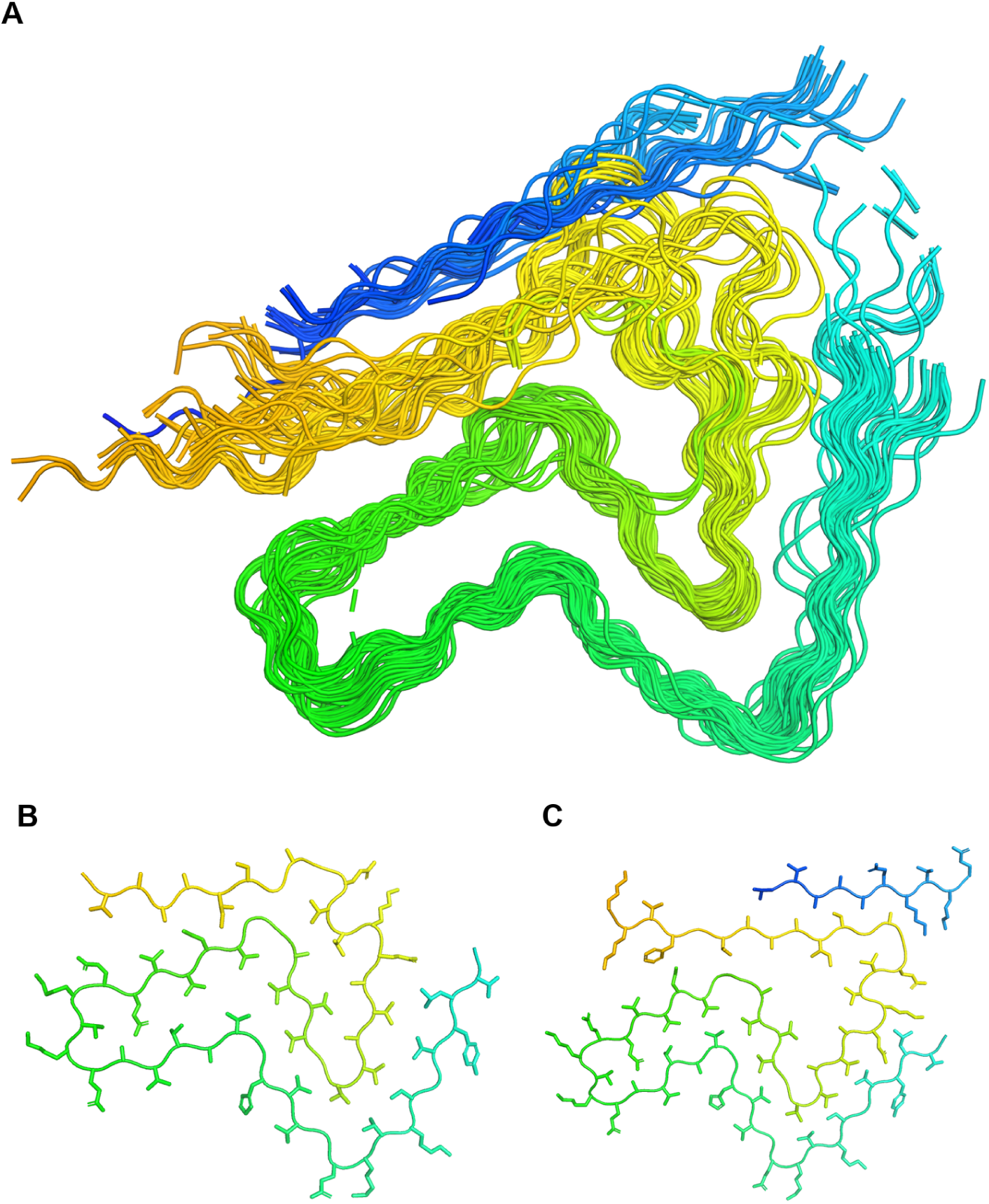
Cluster 2. A) Superposition of all cluster 2 chains, coloured N (blue) to C (red) terminus. B) Medoid 7OZH_A and C) the chain furthest from the medoid, 8CYY_I, shown as single chains with side chains displayed.

**Supplementary Figure 3.**
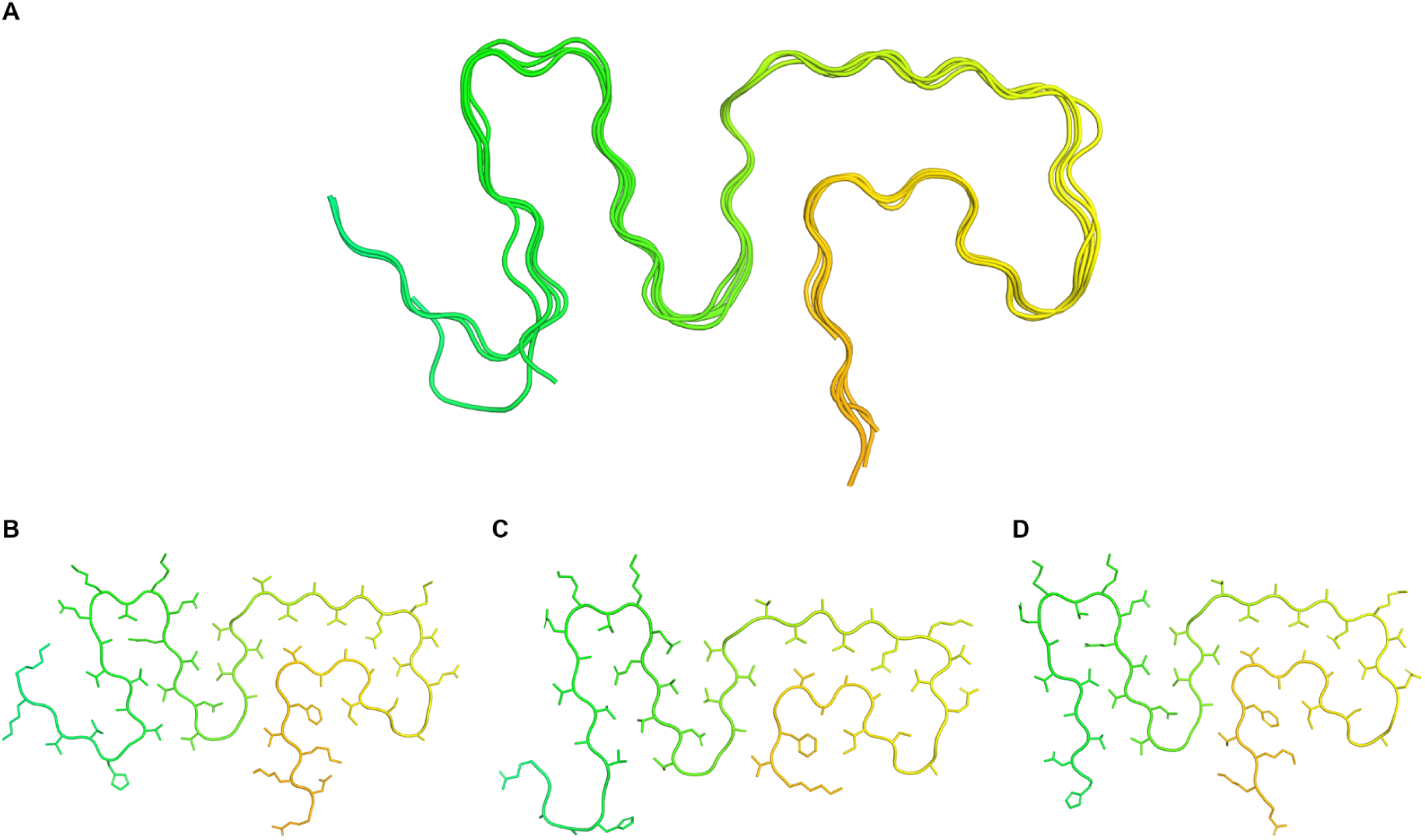
Cluster 3. A) Superposition of all cluster 3 chains, coloured N (blue) to C (red) terminus. B) Medoid 8X7O_A, C) the G51D fibril 7E0F and D) the E46K cross-seeded wild-type fibril 7C1D, shown as single chains with side chains displayed.

**Supplementary Figure 4.**
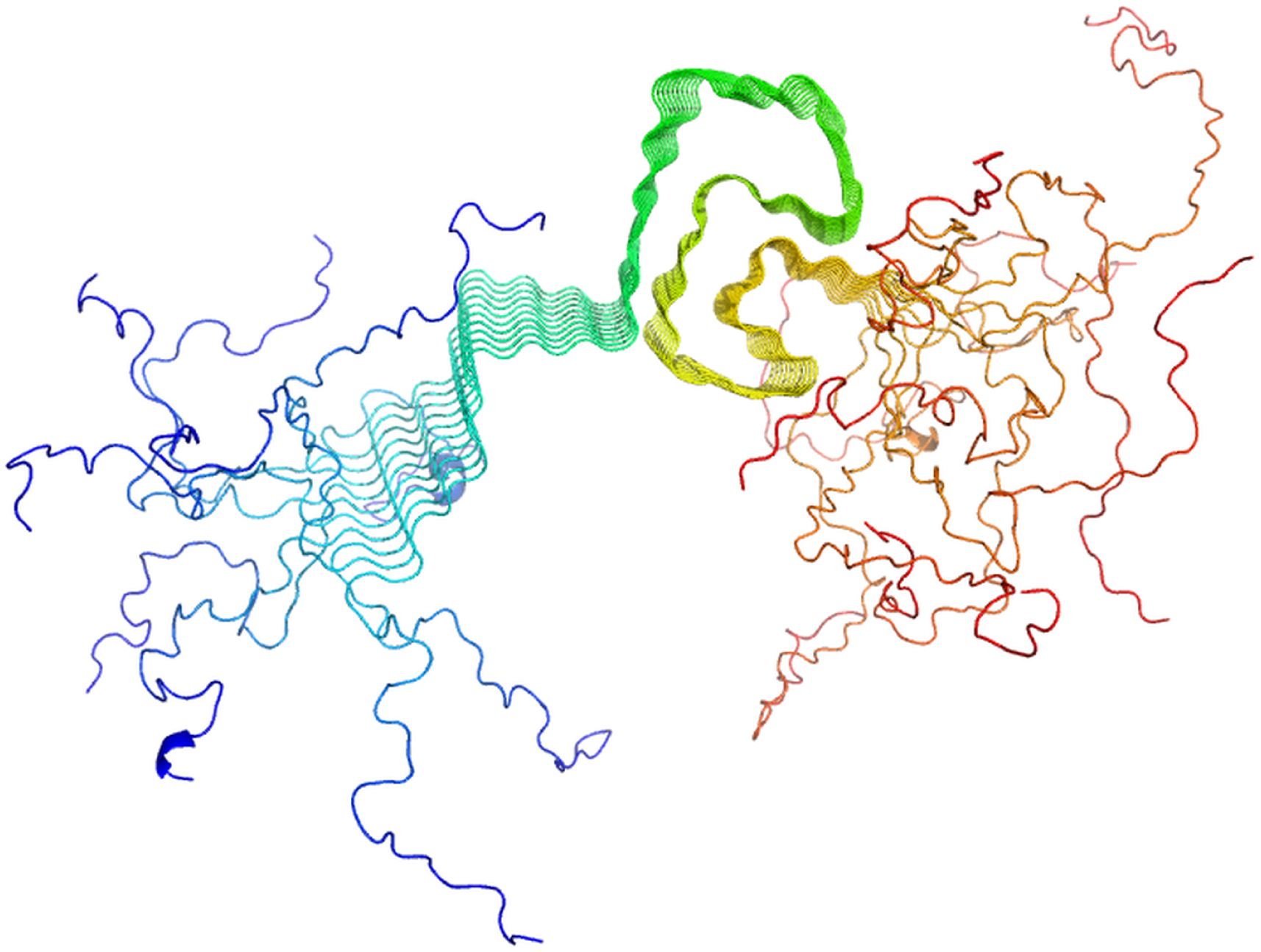
Cluster 4. Superposition of the ten chains of the solid-state NMR structure 2N0A, coloured N (blue) to C (red) terminus.

**Supplementary Figure 5.**
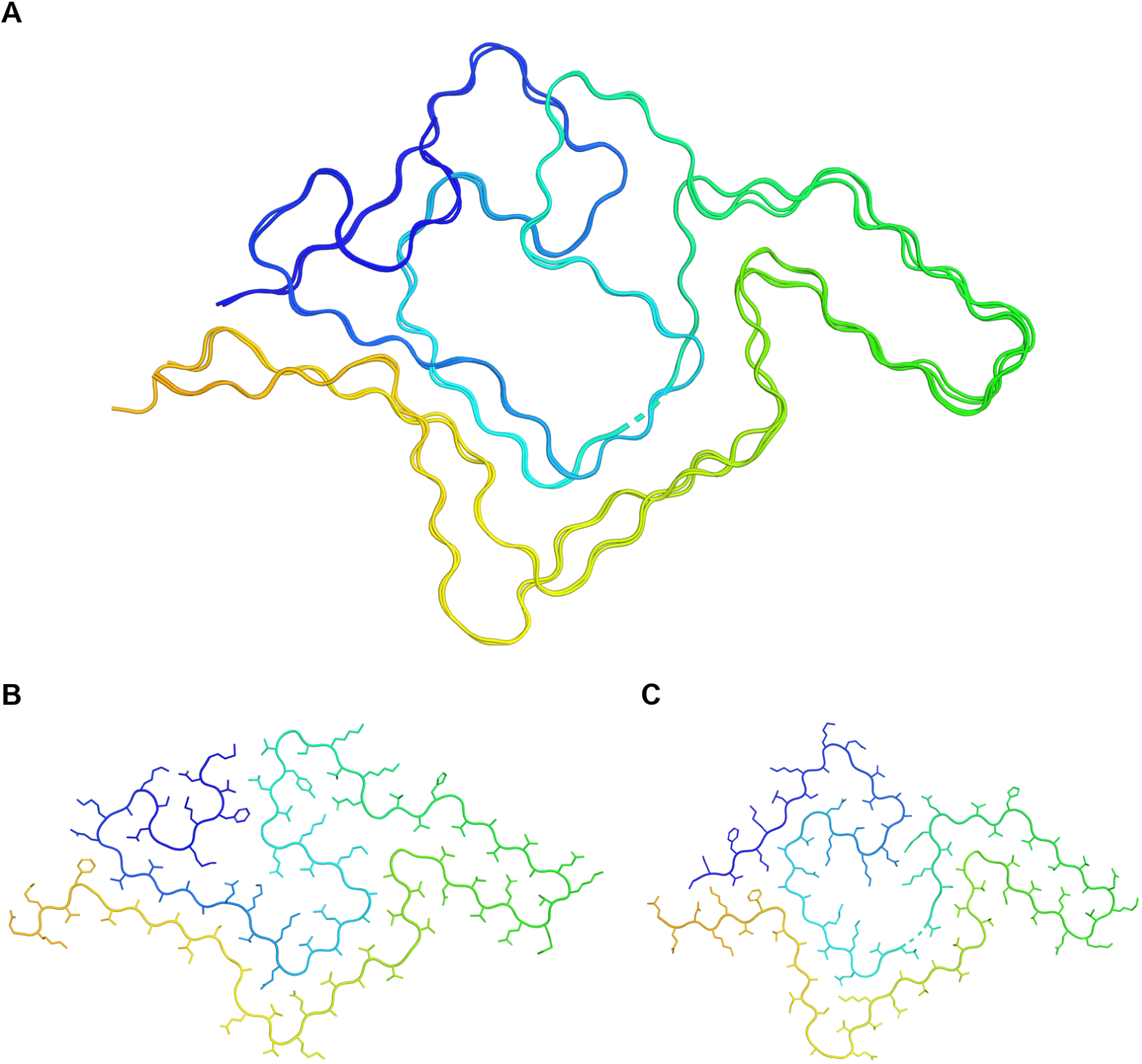
Cluster 5. A) Superposition of all 5 cluster 5 chains. B) The medoid, 8ZLP_A, and C) the chain furthest from the medoid, [6L1T_A] (TM 0.56), shown as single chains with side chains displayed. All structures are coloured N (blue) to C (red) terminus.

**Supplementary Figure 6.**
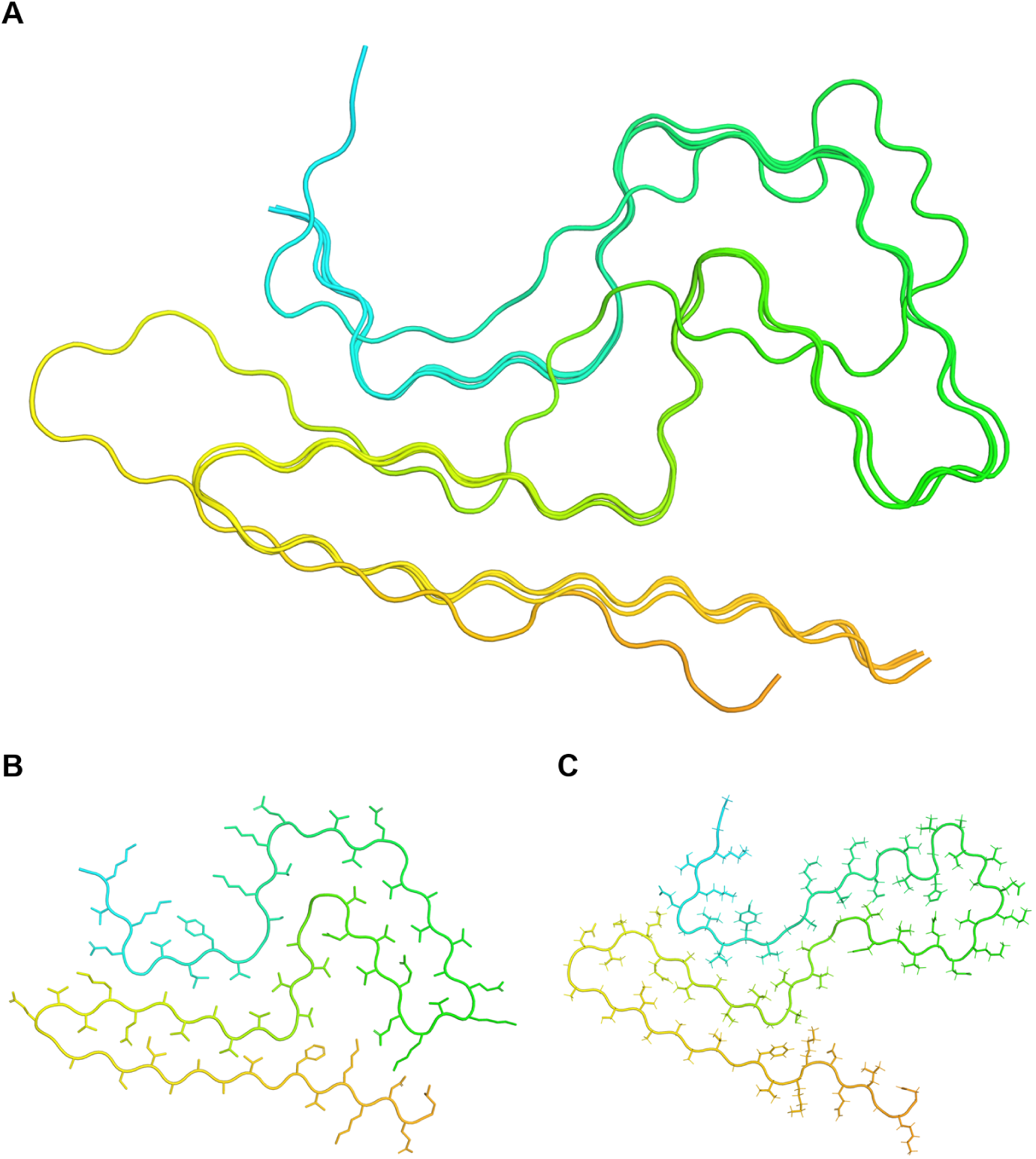
Cluster 6. A) Superposition of all 4 cluster 6 chains. B) The medoid, 9O9K_A, and C) the chain furthest from the medoid, 8FPT (TM 0.54), shown as single chains with side chains displayed. All structures are coloured N (blue) to C (red) terminus.

**Supplementary Figure 7.**
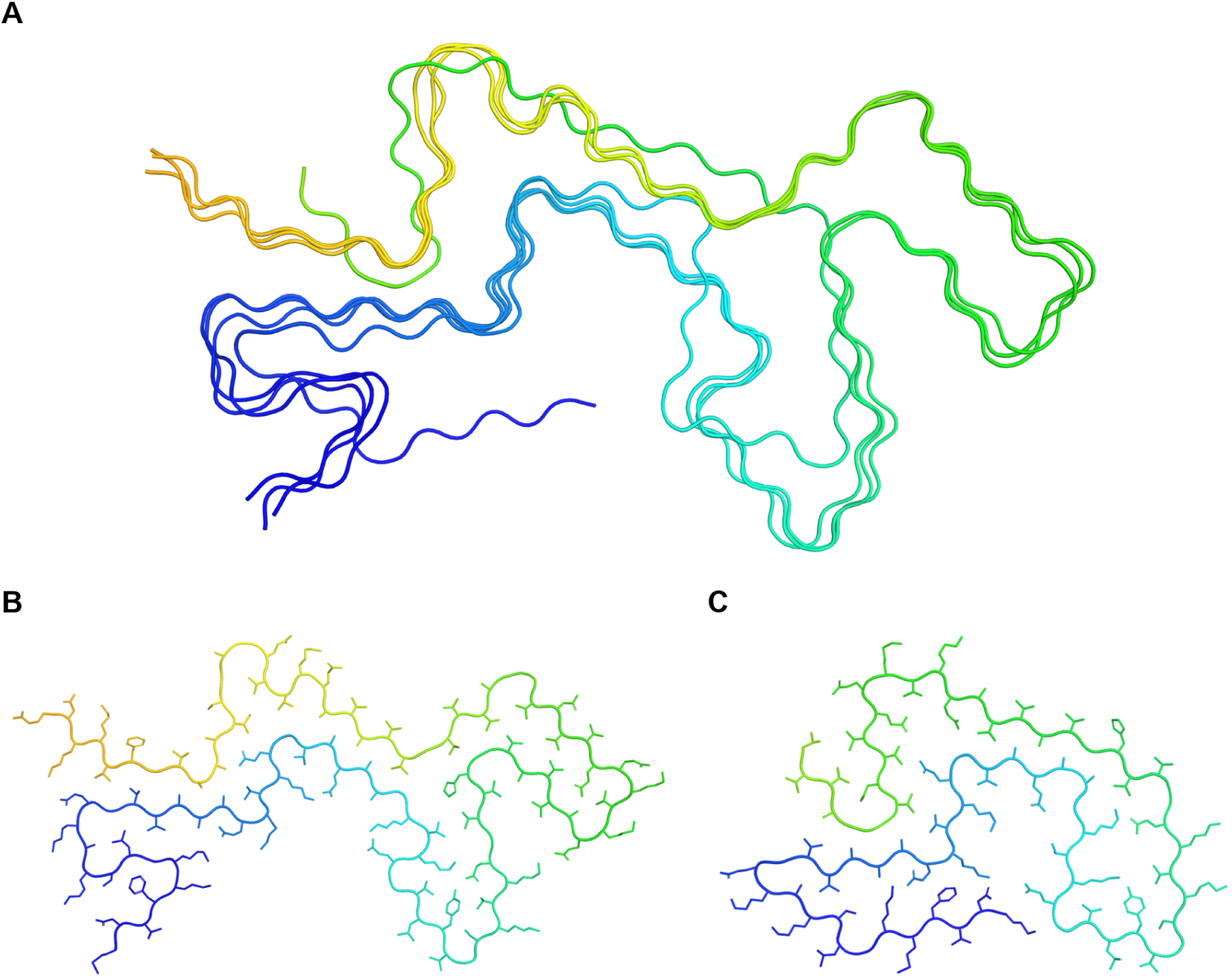
Cluster 7. A) Superposition of all 4 cluster 7 chains. B) The medoid, 8ADU_A, and C) the chain furthest from the medoid, the pSer87 fibril 8JEY (TM 0.52), shown as single chains with side chains displayed. All structures are coloured N (blue) to C (red) terminus.

**Supplementary Figure 8.**
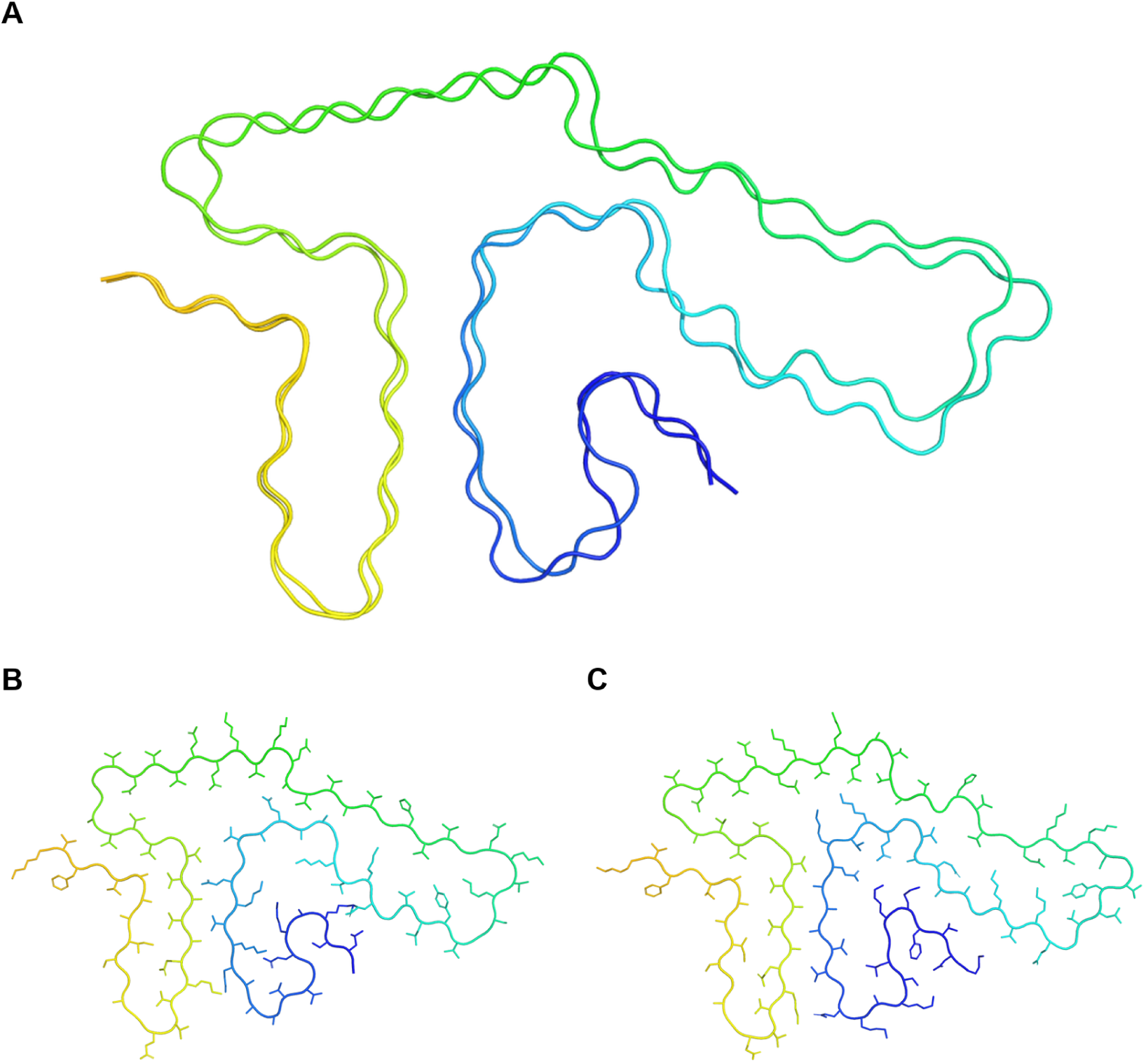
Cluster 8. A) Superposition of both cluster 8 chains. B) The medoid, 8GF7_A, and C) 9D5C (TM 0.87), shown as single chains with side chains displayed. All structures are coloured N (blue) to C (red) terminus.

**Supplementary Figure 9.**
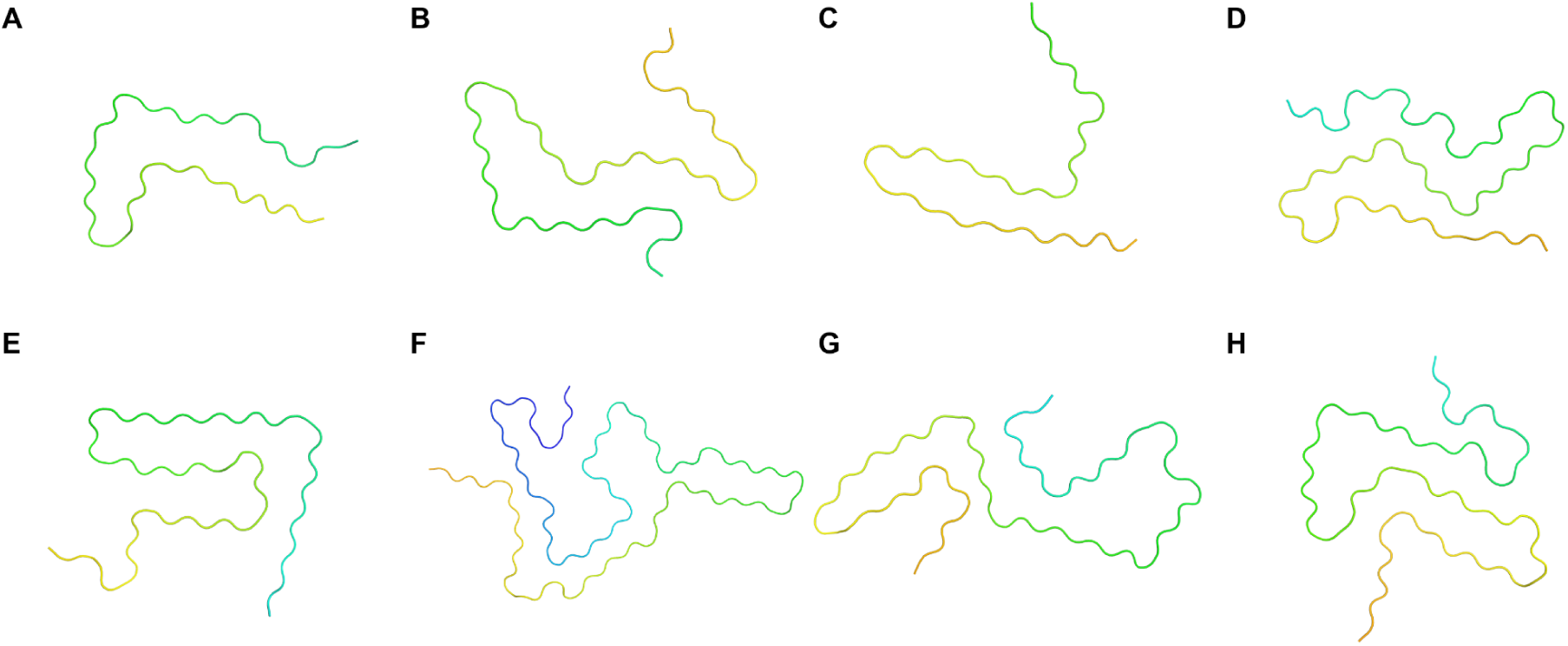
Singleton structures. The eight chains that do not cluster with any other structure at the TM-score threshold, shown as single chains and coloured N (blue) to C (red) terminus. A) 6CU8_A, B) 7LC9_A, C) 7LC9_F, D) 7V4C_A, E) 8JEX_A, F) 9IC7_A, in G) 9W06_B and H) 9W08_D.

**Supplementary Figure 10.**
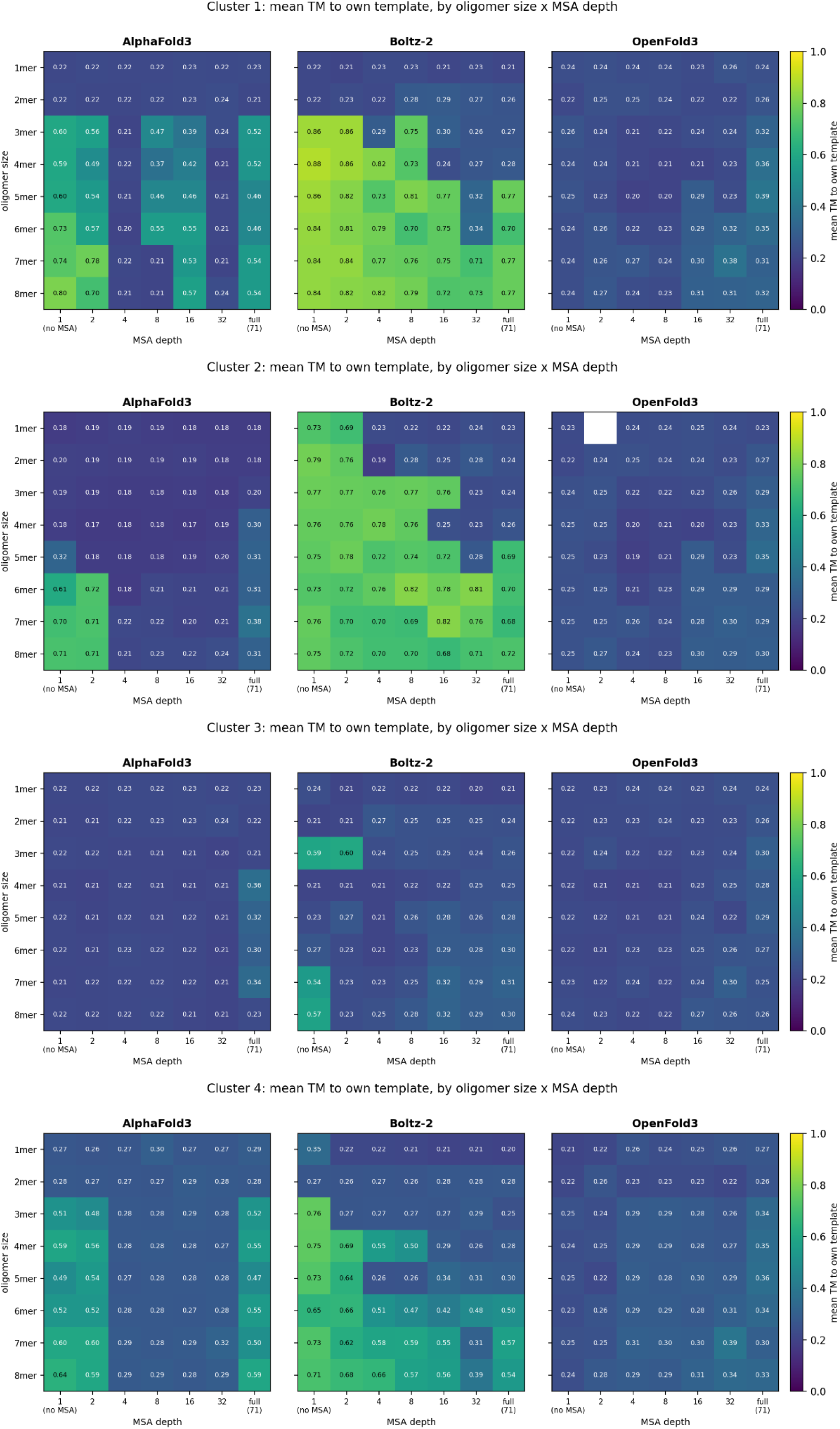
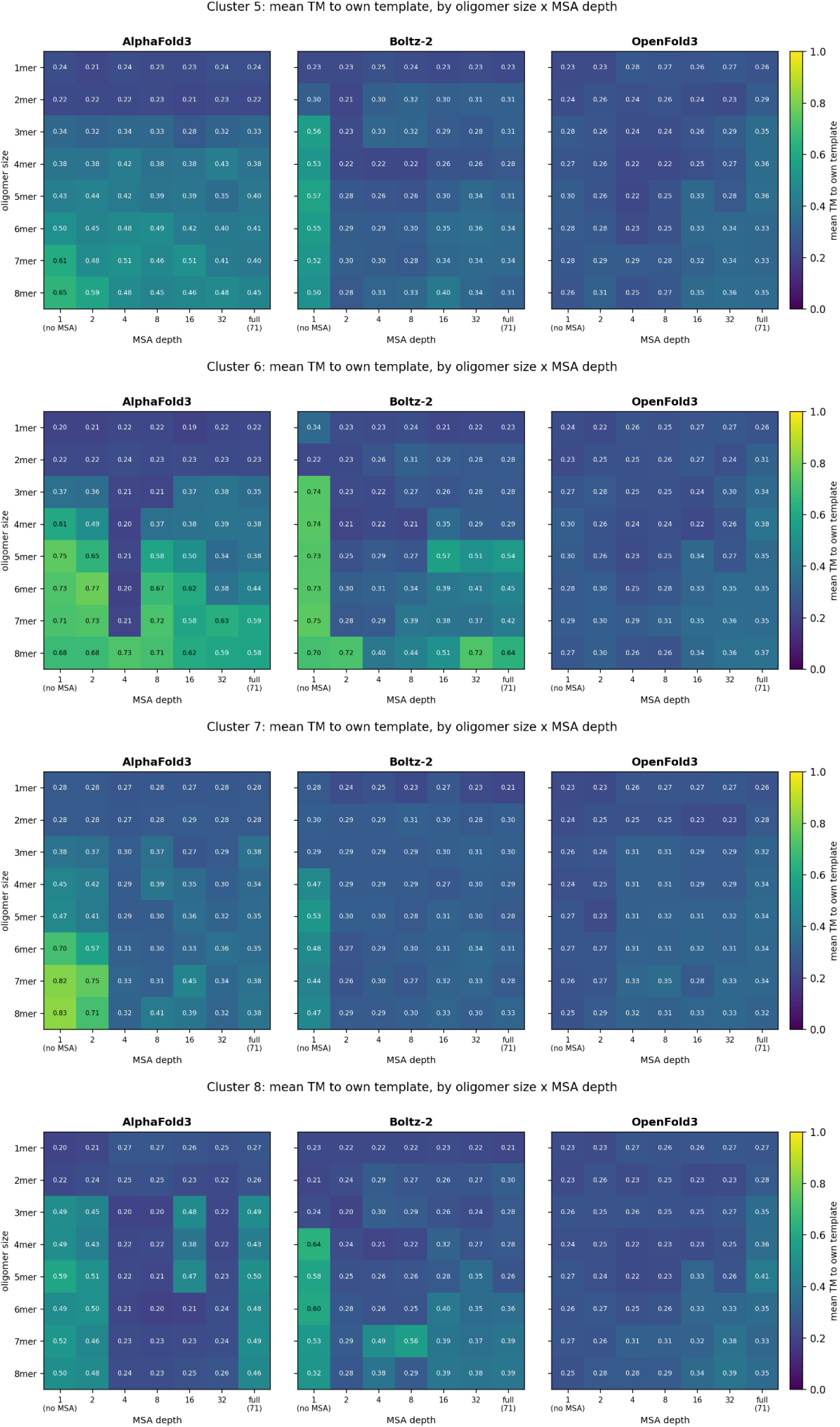
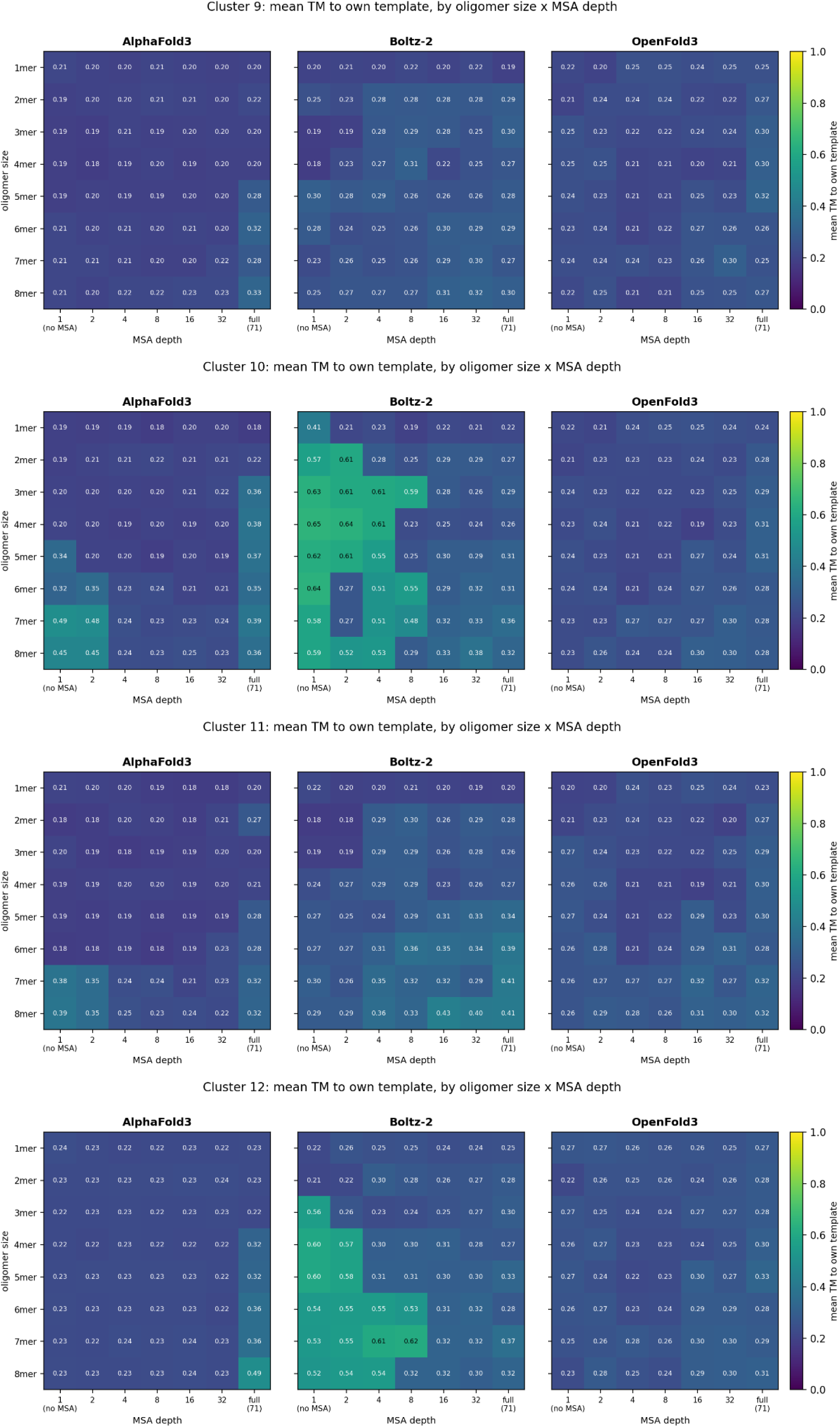
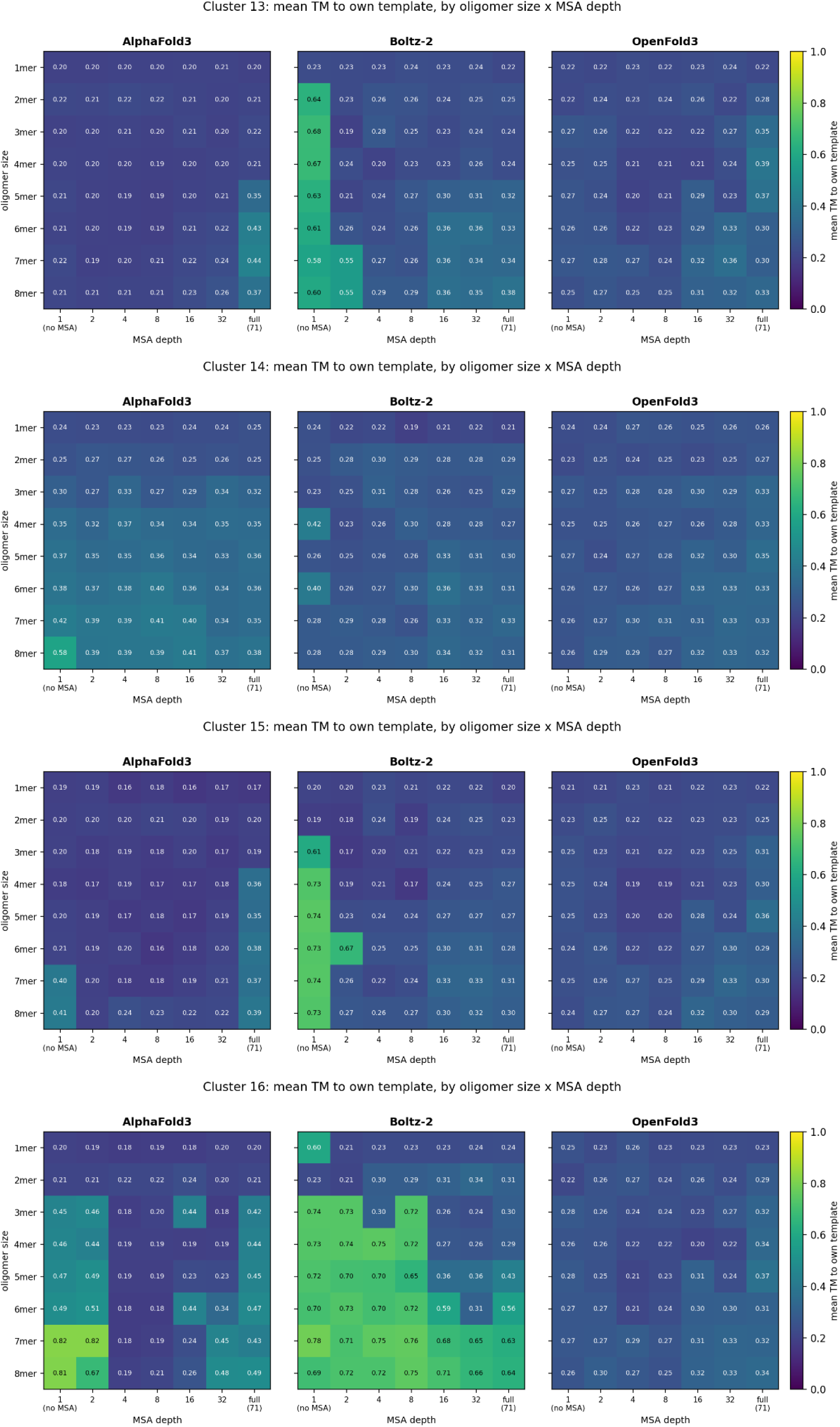
Template obedience by oligomer size and MSA depth. Mean TM-score to the supplied template for each of the 16 target polymorphs, shown by oligomer size (1 to 8 chains) and MSA depth (1, 2, 4, 8, 16, 32 and the full alignment of 71 sequences), for AF3, Boltz-2 and OF3. Each cell is the mean of five models. Targets are the medoid chain of each multi-member cluster and each of the eight singleton structures, numbered as in Table 2.

**Supplementary Figure 11.**
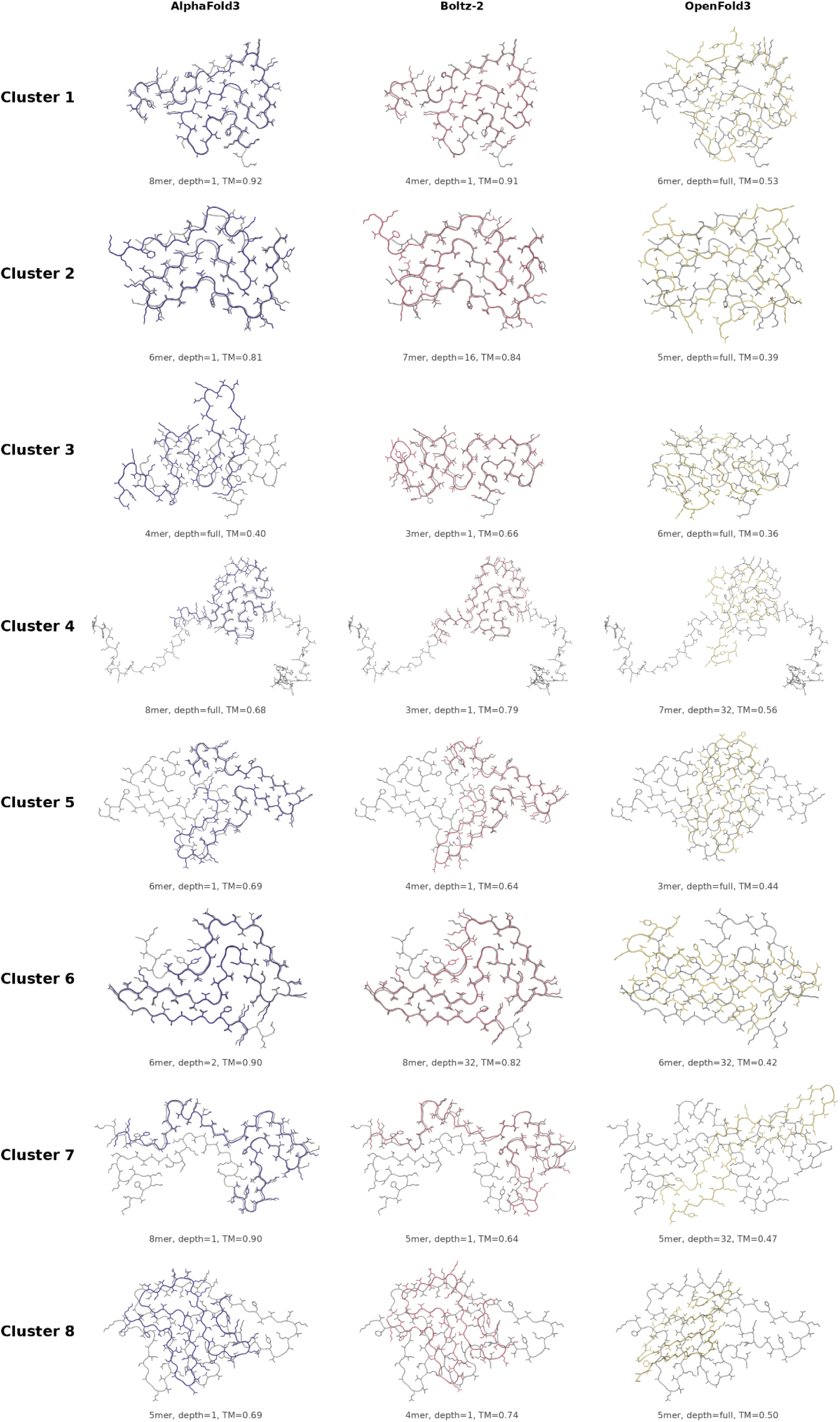
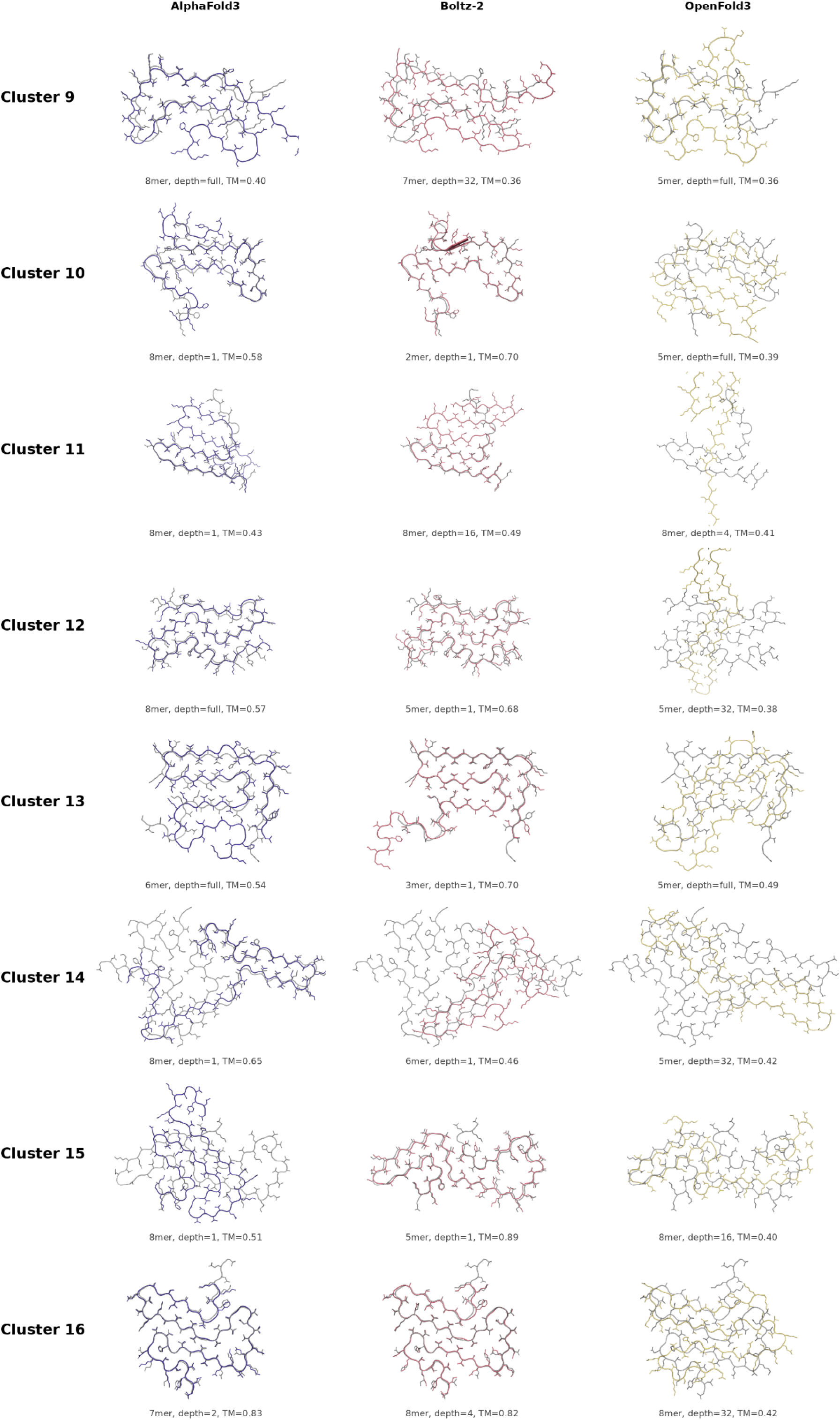
Best-matching templated model for each target polymorph. The highest-scoring model by TM-score to its own template, for each of the 16 targets and each method. The number of chains, MSA depth and TM-score are given for each pair. Template structures are shown in grey, with models coloured by method (AF3 blue, Boltz-2 red, OF3 yellow).

## References

Abramson, J. et al. (2024) ‘Accurate structure prediction of biomolecular interactions with AlphaFold 3’, Nature, 630(8016), pp. 493–500. Available at: 10.1038/s41586-024-07487-w.

Alshammari, M., He, J. and Wriggers, W. (2025) ‘Flexible fitting of AlphaFold2-predicted models to cryo-EM density maps using elastic network models: a methodical affirmation’, Bioinformatics Advances, 5(1), p. vbae181. Available at: 10.1093/bioadv/vbae181.

Aubrey, L.D. and Radford, S.E. (2025) ‘How is the Amyloid Fold Built? Polymorphism and the Microscopic Mechanisms of Fibril Assembly’, Journal of Molecular Biology, 437(22), p. 169008. Available at: 10.1016/j.jmb.2025.169008.

Bæk, K.T. and Kepp, K.P. (2022) ‘Assessment of AlphaFold2 for Human Proteins via Residue Solvent Exposure’, Journal of Chemical Information and Modeling, 62(14), pp. 3391–3400. Available at: 10.1021/acs.jcim.2c00243.

Berman, H., Henrick, K. and Nakamura, H. (2003) ‘Announcing the worldwide Protein Data Bank’, Nature Structural Biology, 10(12), p. 980. Available at: 10.1038/nsb1203-980.

Buel, G.R. and Walters, K.J. (2022) ‘Can AlphaFold2 predict the impact of missense mutations on structure?’, Nature Structural & Molecular Biology, 29(1), pp. 1–2. Available at: 10.1038/s41594-021-00714-2.

Chakravarty, D. et al. (2024) ‘AlphaFold predictions of fold-switched conformations are driven by structure memorization’, Nature Communications, 15(1), p. 7296. Available at: 10.1038/s41467-024-51801-z.

Chiti, F. and Dobson, C.M. (2006) ‘Protein misfolding, functional amyloid, and human disease’, Annual Review of Biochemistry, 75, pp. 333–366. Available at: 10.1146/annurev.biochem.75.101304.123901.

Cock, P.J.A. et al. (2009) ‘Biopython: freely available Python tools for computational molecular biology and bioinformatics’, Bioinformatics, 25(11), pp. 1422–1423. Available at: 10.1093/bioinformatics/btp163.

Cohlberg, J.A. et al. (2002) ‘Heparin and other glycosaminoglycans stimulate the formation of amyloid fibrils from alpha-synuclein in vitro’, Biochemistry, 41(5), pp. 1502–1511. Available at: 10.1021/bi011711s.

Connor, J.P., Radford, S.E. and Brockwell, D.J. (2025) ‘Structural and thermodynamic classification of amyloid polymorphs’, Structure, 33(10), pp. 1793–1804.e3. Available at: 10.1016/j.str.2025.07.005.

Eisenberg, D.S. and Sawaya, M.R. (2017) ‘Structural Studies of Amyloid Proteins at the Molecular Level’, Annual Review of Biochemistry, 86, pp. 69–95. Available at: 10.1146/annurev-biochem-061516-045104.

Elliott, L.G., Simpkin, A.J. and Rigden, D.J. (2025) ‘ABCFold: easier running and comparison of AlphaFold 3, Boltz-1, and Chai-1’, Bioinformatics Advances, 5(1), p. vbaf153. Available at: 10.1093/bioadv/vbaf153.

Fitzpatrick, A.W.P. et al. (2017) ‘Cryo-EM structures of tau filaments from Alzheimer’s disease’, Nature, 547(7662), pp. 185–190. Available at: 10.1038/nature23002.

Frey, L. et al. (2024) ‘On the pH-dependence of α-synuclein amyloid polymorphism and the role of secondary nucleation in seed-based amyloid propagation’, eLife, 12, p. RP93562. Available at: 10.7554/eLife.93562.

Frieg, B. et al. (2022) ‘The 3D structure of lipidic fibrils of α-synuclein’, Nature Communications, 13(1), p. 6810. Available at: 10.1038/s41467-022-34552-7.

Gomez-Gutierrez, R. et al. (2023) ‘Two structurally defined Aβ polymorphs promote different pathological changes in susceptible mice’, EMBO reports, 24(8), p. e57003. Available at: 10.15252/embr.202357003.

Greenwald, J. and Riek, R. (2010) ‘Biology of Amyloid: Structure, Function, and Regulation’, Structure, 18(10), pp. 1244–1260. Available at: 10.1016/j.str.2010.08.009.

Gremer, L. et al. (2017) ‘Fibril structure of amyloid-β(1-42) by cryo-electron microscopy’, *Science (New York*, N.Y.), 358(6359), pp. 116–119. Available at: 10.1126/science.aao2825.

Guerrero-Ferreira, R. et al. (2018) ‘Cryo-EM structure of alpha-synuclein fibrils’, eLife, 7, p. e36402. Available at: 10.7554/eLife.36402.

He, S. and Scheres, S.H.W. (2017a) ‘Helical reconstruction in RELION’, Journal of Structural Biology, 198(3), pp. 163–176. Available at: 10.1016/j.jsb.2017.02.003.

He, S. and Scheres, S.H.W. (2017b) ‘Helical reconstruction in RELION’, Journal of Structural Biology, 198(3), pp. 163–176. Available at: 10.1016/j.jsb.2017.02.003.

He, Y., Yu, Z. and Chen, S. (2019) ‘Alpha-Synuclein Nitration and Its Implications in Parkinson’s Disease’, ACS chemical neuroscience, 10(2), pp. 777–782. Available at: 10.1021/acschemneuro.8b00288.

Hu, J. et al. (2024) ‘Phosphorylation and O-GlcNAcylation at the same α-synuclein site generate distinct fibril structures’, Nature Communications, 15(1), p. 2677. Available at: 10.1038/s41467-024-46898-1.

Jumper, J. et al. (2021) ‘Highly accurate protein structure prediction with AlphaFold’, Nature, 596(7873), pp. 583–589. Available at: 10.1038/s41586-021-03819-2.

Kabsch, W. (1976) ‘A solution for the best rotation to relate two sets of vectors’, Acta Crystallographica Section A, 32(5), pp. 922–923. Available at: 10.1107/S0567739476001873.

Kabsch, W. and Sander, C. (1983) ‘Dictionary of protein secondary structure: pattern recognition of hydrogen-bonded and geometrical features’, Biopolymers, 22(12), pp. 2577–2637. Available at: 10.1002/bip.360221211.

van der Kant, R. et al. (2022) ‘Thermodynamic analysis of amyloid fibril structures reveals a common framework for stability in amyloid polymorphs’, Structure, 30(8), pp. 1178–1189.e3. Available at: 10.1016/j.str.2022.05.002.

van Kempen, M. et al. (2024) ‘Fast and accurate protein structure search with Foldseek’, Nature Biotechnology, 42(2), pp. 243–246. Available at: 10.1038/s41587-023-01773-0.

Kendrew, J.C. et al. (1958) ‘A Three-Dimensional Model of the Myoglobin Molecule Obtained by X-Ray Analysis’, Nature, 181(4610), pp. 662–666. Available at: 10.1038/181662a0.

Kochen, N.N. et al. (2024) ‘Post-translational modification sites are present in hydrophilic cavities of alpha-synuclein, tau, FUS, and TDP-43 fibrils: A molecular dynamics study’, Proteins, 92(7), pp. 854–864. Available at: 10.1002/prot.26679.

LaFerla, F.M., Green, K.N. and Oddo, S. (2007) ‘Intracellular amyloid-β in Alzheimer’s disease’, Nature Reviews Neuroscience, 8(7), pp. 499–509. Available at: 10.1038/nrn2168.

Li, B. et al. (2018a) ‘Cryo-EM of full-length α-synuclein reveals fibril polymorphs with a common structural kernel’, Nature Communications, 9(1), p. 3609. Available at: 10.1038/s41467-018-05971-2.

Li, B. et al. (2018b) ‘Cryo-EM of full-length α-synuclein reveals fibril polymorphs with a common structural kernel’, Nature Communications, 9(1), p. 3609. Available at: 10.1038/s41467-018-05971-2.

Li, D. and Liu, C. (2023) ‘Molecular rules governing the structural polymorphism of amyloid fibrils in neurodegenerative diseases’, Structure, 31(11), pp. 1335–1347. Available at: 10.1016/j.str.2023.08.006.

Li, X. et al. (2013) ‘Electron counting and beam-induced motion correction enable near-atomic-resolution single-particle cryo-EM’, Nature Methods, 10(6), pp. 584–590. Available at: 10.1038/nmeth.2472.

Long, H. et al. (2021) ‘Wild-type α-synuclein inherits the structure and exacerbated neuropathology of E46K mutant fibril strain by cross-seeding’, Proceedings of the National Academy of Sciences of the United States of America, 118(20), p. e2012435118. Available at: 10.1073/pnas.2012435118.

Lövestam, S. et al. (2021) ‘Seeded assembly in vitro does not replicate the structures of α-synuclein filaments from multiple system atrophy’, FEBS open bio, 11(4), pp. 999–1013. Available at: 10.1002/2211-5463.13110.

Lutter, L., Aubrey, L.D. and Xue, W.-F. (2021) ‘On the Structural Diversity and Individuality of Polymorphic Amyloid Protein Assemblies’, Journal of Molecular Biology, 433(20), p. 167124. Available at: 10.1016/j.jmb.2021.167124.

Mesdaghi, S. et al. (2023) ‘Deep Learning-based structure modelling illuminates structure and function in uncharted regions of β-solenoid fold space’, Journal of Structural Biology, 215(3), p. 108010. Available at: 10.1016/j.jsb.2023.108010.

Milchberg, M.H. et al. (2025) ‘Alpha-Synuclein Fibril Structures Cluster into Distinct Classes’. bioRxiv: The Preprint Server for Biology. Available at: 10.1101/2025.04.30.651534.

Millán, C. et al. (2023) ‘Likelihood-based docking of models into cryo-EM maps’, *Acta Crystallographica. Section D*, Structural Biology, 79(Pt 4), pp. 281–289. Available at: 10.1107/S2059798323001602.

Mirdita, M. et al. (2022) ‘ColabFold: making protein folding accessible to all’, Nature Methods, 19(6), pp. 679–682. Available at: 10.1038/s41592-022-01488-1.

Nelson, R. et al. (2005) ‘Structure of the cross-beta spine of amyloid-like fibrils’, Nature, 435(7043), pp. 773–778. Available at: 10.1038/nature03680.

Ohgita, T. et al. (2025) ‘Kinetic Mechanism of Heparin-Induced Fibrillation of α-Synuclein’, ACS chemical neuroscience, 16(21), pp. 4246–4256. Available at: 10.1021/acschemneuro.5c00489.

Otzen, D. and Riek, R. (2019) ‘Functional Amyloids’, Cold Spring Harbor Perspectives in Biology, 11(12), p. a033860. Available at: 10.1101/cshperspect.a033860.

Pak, M.A. et al. (2023) ‘Using AlphaFold to predict the impact of single mutations on protein stability and function’, PLOS ONE, 18(3), p. e0282689. Available at: 10.1371/journal.pone.0282689.

Paravastu, A.K. et al. (2008) ‘Molecular structural basis for polymorphism in Alzheimer’s β-amyloid fibrils’, Proceedings of the National Academy of Sciences, 105(47), pp. 18349–18354. Available at: 10.1073/pnas.0806270105.

Passaro, S. et al. (2025) ‘Boltz-2: Towards Accurate and Efficient Binding Affinity Prediction’. bioRxiv: The Preprint Server for Biology. Available at: 10.1101/2025.06.14.659707.

Pedregosa, F. et al. (2011) ‘Scikit-learn: Machine Learning in Python’, The Journal of Machine Learning Research, 12(null), pp. 2825–2830.

Perutz, M.F. et al. (1960) ‘Structure of Hæmoglobin: A Three-Dimensional Fourier Synthesis at 5.5-Å. Resolution, Obtained by X-Ray Analysis’, Nature, 185(4711), pp. 416–422. Available at: 10.1038/185416a0.

Petkova, A.T. et al. (2002) ‘A structural model for Alzheimer’s β-amyloid fibrils based on experimental constraints from solid state NMR’, Proceedings of the National Academy of Sciences, 99(26), pp. 16742–16747. Available at: 10.1073/pnas.262663499.

Pinheiro, F., Santos, J. and Ventura, S. (2021) ‘AlphaFold and the amyloid landscape’, Journal of Molecular Biology, 433(20), p. 167059. Available at: 10.1016/j.jmb.2021.167059.

Pradhan, B. et al. (2026) ‘Atomic structures of medin and Aβ fibrils reveal polymorphic remodeling in mixed amyloid systems’, Nature Communications, 17(1), p. 5804. Available at: 10.1038/s41467-026-72515-4.

Pratt, O.S. et al. (2025) ‘AlphaFold 2, but not AlphaFold 3, predicts confident but unrealistic β-solenoid structures for repeat proteins’, Computational and Structural Biotechnology Journal, 27, pp. 467–477. Available at: 10.1016/j.csbj.2025.01.016.

Radamaker, L. et al. (2019) ‘Cryo-EM structure of a light chain-derived amyloid fibril from a patient with systemic AL amyloidosis’, Nature Communications, 10(1), p. 1103. Available at: 10.1038/s41467-019-09032-0.

Rambaran, R.N. and Serpell, L.C. (2008) ‘Amyloid fibrils’, Prion, 2(3), pp. 112–117.

Sanchez, J.C. et al. (2024) ‘High-Resolution Cryo-EM Structure Determination of α-synuclein - A Prototypical Amyloid Fibril’. bioRxiv: The Preprint Server for Biology. Available at: 10.1101/2024.09.18.613698.

Sawaya, M.R. et al. (2007a) ‘Atomic structures of amyloid cross-β spines reveal varied steric zippers’, Nature, 447(7143), pp. 453–457. Available at: 10.1038/nature05695.

Sawaya, M.R. et al. (2007b) ‘Atomic structures of amyloid cross-β spines reveal varied steric zippers’, Nature, 447(7143), pp. 453–457. Available at: 10.1038/nature05695.

Sawaya, M.R. et al. (2021) ‘The expanding amyloid family: Structure, stability, function, and pathogenesis’, Cell, 184(19), pp. 4857–4873. Available at: 10.1016/j.cell.2021.08.013.

Scheres, S.H.W. (2026) ‘The amyloid packing difference: a pairwise comparison metric for amyloid structures’. bioRxiv, p. 2026.02.18.706523. Available at: 10.64898/2026.02.18.706523.

Scheres, S.H.W., Ryskeldi-Falcon, B. and Goedert, M. (2023) ‘Molecular pathology of neurodegenerative diseases by cryo-EM of amyloids’, Nature, 621(7980), pp. 701–710. Available at: 10.1038/s41586-023-06437-2.

Schweighauser, M. et al. (2020) ‘Structures of α-synuclein filaments from multiple system atrophy’, Nature, 585(7825), pp. 464–469. Available at: 10.1038/s41586-020-2317-6.

Shahnawaz, M. et al. (2020) ‘Discriminating α-synuclein strains in Parkinson’s disease and multiple system atrophy’, Nature, 578(7794), pp. 273–277. Available at: 10.1038/s41586-020-1984-7.

Sønderby, T.V. et al. (2022) ‘Functional Bacterial Amyloids: Understanding Fibrillation, Regulating Biofilm Fibril Formation and Organizing Surface Assemblies’, Molecules, 27(13). Available at: 10.3390/molecules27134080.

Steinegger, M. and Söding, J. (2017) ‘MMseqs2 enables sensitive protein sequence searching for the analysis of massive data sets’, Nature Biotechnology, 35(11), pp. 1026–1028. Available at: 10.1038/nbt.3988.

Taylor, A.I.P. and Staniforth, R.A. (2022) ‘General Principles Underpinning Amyloid Structure’, Frontiers in Neuroscience, 16. Available at: 10.3389/fnins.2022.878869.

The OpenFold3 Team (2026) ‘OpenFold3-preview’. Available at: https://github.com/aqlaboratory/openfold-3 (Accessed: 12 October 2026).

Tian, P. et al. (2015) ‘Structure of a Functional Amyloid Protein Subunit Computed Using Sequence Variation’, Journal of the American Chemical Society, 137(1), pp. 22–25. Available at: 10.1021/ja5093634.

Tycko, R. (2015) ‘Amyloid polymorphism: structural basis and neurobiological relevance’, Neuron, 86(3), pp. 632–645. Available at: 10.1016/j.neuron.2015.03.017.

Verma, M., Vats, A. and Taneja, V. (2015) ‘Toxic species in amyloid disorders: Oligomers or mature fibrils’, Annals of Indian Academy of Neurology, 18(2), pp. 138–145. Available at: 10.4103/0972-2327.144284.

Virtanen, P. et al. (2020) ‘SciPy 1.0: fundamental algorithms for scientific computing in Python’, Nature Methods, 17(3), pp. 261–272. Available at: 10.1038/s41592-019-0686-2.

Watts, J.C. (2025) ‘Shedding light on the α-synuclein fibril fuzzy coat’, Neuron, 113(11), pp. 1653–1655. Available at: 10.1016/j.neuron.2025.04.022.

Wells, C. et al. (2021) ‘The role of amyloid oligomers in neurodegenerative pathologies’, International Journal of Biological Macromolecules, 181, pp. 582–604. Available at: 10.1016/j.ijbiomac.2021.03.113.

Xu, J. and Zhang, Y. (2010) ‘How significant is a protein structure similarity with TM-score = 0.5?’, Bioinformatics, 26(7), pp. 889–895. Available at: 10.1093/bioinformatics/btq066.

Yang, Y. et al. (2022) ‘Structures of α-synuclein filaments from human brains with Lewy pathology’, Nature, 610(7933), pp. 791–795. Available at: 10.1038/s41586-022-05319-3.

Yang, Y. et al. (2023) ‘New SNCA mutation and structures of α-synuclein filaments from juvenile-onset synucleinopathy’, Acta Neuropathologica, 145(5), pp. 561–572. Available at: 10.1007/s00401-023-02550-8.

Zhang, Y. and Skolnick, J. (2005) ‘TM-align: a protein structure alignment algorithm based on the TM-score’, Nucleic Acids Research, 33(7), pp. 2302–2309. Available at: 10.1093/nar/gki524.

Zhao, Y. et al. (2023) ‘Metal Exposure and Risk of Parkinson Disease: A Systematic Review and Meta-Analysis’, American Journal of Epidemiology, 192(7), pp. 1207–1223. Available at: 10.1093/aje/kwad082.

Zielinski, M., Röder, C. and Schröder, G.F. (2021) ‘Challenges in sample preparation and structure determination of amyloids by cryo-EM’, The Journal of Biological Chemistry, 297(2), p. 100938. Available at: 10.1016/j.jbc.2021.100938.

